# GP130 Y814 SIGNALING IS REQUIRED FOR THE DYNAMIN-MEDIATED ENDOCYTOSIS, MAPK/P38 ACTIVATION AND PERSISTENCE OF CHRONIC SYSTEMIC INFLAMMATION INDUCED BY HIGH FAT DIET

**DOI:** 10.64898/2026.02.06.704505

**Authors:** Jade Tassey, Youngjoo Lee, Arijita Sarkar, Jichen Yang, Siyoung Lee, Una Stevic, Nancy Q. Liu, Jinxiu Lu, Andrew C. Drake, Vanessa Jabbour, Mary Vergel, Ana C. Maretti-Mira, Lucy Golden-Mason, Rong Lu, Denis Evseenko

**Author notes:** Correspondence to: Denis Evseenko MD, PhD. Professor of Orthopaedic Surgery, Stem Cell Research and Regenerative Medicine. USC, Los Angeles, CA, 90033, USA. These authors contributed equally to this study (Jade Tassey, Youngjoo Lee).

## Abstract

In chronic inflammatory diseases such as obesity, tissue and cellular homeostasis are disrupted by persistent low-grade inflammation, where one of the most prominent pro-inflammatory cytokines that correlates with age and body mass index (BMI) is IL-6. All members of the IL-6 family of cytokines signal through their obligate co-receptor gp130, in whichsignaling tyrosines on the intracellular portion of gp130 activate multiple downstream pathways such as the canonical STAT and MAPK pathways. However, non-canonical gp130 pathways such as SRC family of kinases (SFKs) signaling have emerged as drivers of cellular stress response. Our recently published mutant mouse model carrying a constitutive inactivation of gp130-Y814 (F814 mice), which impairs SFK activation, exhibited an enhanced resolution of inflammatory responses and improved regenerative outcomes in both acute skin wound healing and post-traumatic osteoarthritis models. The current study was designed to explore whether the gp130-Y814 mutation reduces systemic chronic inflammation and multimorbidity in a high-fat diet (HFD)-induced model and to interrogate the possible downstream cellular mechanisms that are affected. In response to HFD, F814 mice showed significantly reduced systemic and tissue-specific inflammatory responses and protection from obesity-induced bone loss and osteoarthritis compared to wild type (WT) mice. After extensive characterization, the role of gp130-Y814 in monocytes/macrophages appeared to be dispensable, but we discovered that the F814 mutation blunts gp130 receptor internalization, p38/MAPK activation and the release of matrix metalloproteinases (MMPs) in chondrocytes. Finally, we showed that F814 chondrocytes had markedly reduced activation of the SFK-dependent GTPase dynamin 2 (Dyn2) in response to catabolic IL-6 cytokines. Introduction of a dominant-negative Dyn2 mutant into WT chondrocytes phenocopied the effects of the F814 mutation, resulting in attenuated p38/MAPK signaling and reduced MMP activation following stimulation with IL-6 cytokines. This study demonstrates, for the first time, that the Y814 residue is directly implicated in Dyn2-mediated internalization of the gp130 receptor, thereby modulating downstream signaling and contributing to pathological outcomes in chronic inflammatory and degenerative diseases in a cell type–specific manner.

## Introduction

Inflammation refers to the body’s natural response to injury or infection; without it, organisms would be unable to combat harmful stimuli or initiate tissue repair (1). Differing from acute inflammation, chronic inflammation is defined as the persistence of low-grade, sterile systemic inflammation characterized by the continual activation of immune and surrounding mesenchymal cells (2). This shift in inflammatory response can cause a breakdown of immune tolerance and lead to major alterations in all organs, linking chronic inflammation to a variety of diseases such as cancer, cardiovascular and liver disorders, diabetes, and arthritis (3).

Several factors such as aging, diet, obesity, and stress can promote chronic inflammation, in which biomarkers like C-reactive protein (CRP), Monocyte Chemoattractant Protein-1 (MCP-1), and pro-inflammatory cytokines such as Tumor necrosis factor (TNFα), Interleukin (IL)-1β, and IL-6 are greatly heightened (4,5). Obesity-induced osteoarthritis (OA) is caused by the overloading of joints due to excessive weight and systemic inflammation, ultimately leading to the destruction of articular cartilage (6). While the activity of pro-inflammatory myeloid cells such as monocytes/macrophages in chronic inflammatory diseases such as OA has been described (6), chondrocytes (the cells that synthesize and maintain cartilage matrix) begin excessively producing matrix metalloproteinases (MMPs) that break down the collagens and proteoglycans of cartilage (7).

IL-6 is traditionally accepted as one of the top three major pro-inflammatory cytokines (the others being IL-1β and TNFα), where increasing IL-6 levels have been directly correlated with increased age and body mass index (8,9). IL-6 is just one of the several members of IL-6 family of cytokines, which also include Oncostatin M (OSM), IL-11, and Leukemia Inhibitory Factor (LIF). All IL-6 family cytokines signal through the obligate membrane-bound co-receptor glycoprotein 130 (gp130; *Il6st*) to activate downstream intracellular pathways (10). Activation of intracellular gp130-associated Janus kinases (JAKs) results in the phosphorylation of tyrosine (Y) residues (Y759, Y767, Y814, Y905, and Y915) within the cytoplasmic domain of gp130 (11). These phosphorylated tyrosine residues serve as docking sites for multiple downstream signaling pathways, including JAK/STAT, Ras/ERK1/2/MAPK, and the PI3K/AKT cascades (12). However, there has been growing interest in non-canonical gp130 signaling pathways, including SRC Family of Kinases (SFKs) and p38/MAPK pathways (13,14).

Recent studies suggest that receptor complexes like gp130 are internalized into endosomes and can serve as signaling platforms that sustain intracellular signaling, potentially with altered signaling quality (15,16). In this context, it is plausible that endocytosed gp130 can sustain signaling and engage downstream pathways that differ in strength and nature from those initiated at the plasma membrane. For example, STAT3 signaling can be activated and sustained at the plasma membrane, whereas ERK1/2/MAPK signaling is dependent on endocytosis (17). Other studies suggest that internalization of gp130 is p38/MAPK dependent (18), adding to the complexities of gp130 endocytic signaling. SFKs have been shown to directly phosphorylate dynamin, a GTPase crucial for endocytic vesicle scission from the plasma membrane and the Golgi body (19), and co-localization of SFKs in endosomes (both early and late endosomes) has been widely reported (20), even in the context of OA (21). SFK-p38/MAPK interplay has also been described (22), suggesting an interesting link between the two signaling pathways and endocytosed receptors such as gp130. SFKs also phosphorylate the inhibitor of Rho GTPases (Rho GDI), which allows Rho GTPases to become activated and contribute to essential functions for membrane trafficking such as migration and proliferation (23). Collectively, these multiple downstream targets of gp130/SFK endosomal signaling may appear as independent mechanisms, but it could be argued that they are interconnected . A deeper understanding of how these pathways converge and orchestrate cellular responses could reveal a new paradigm of potential biological targets for investigation.

We have previously shown that gp130-Y814 activates SFKs, and that the mutation of Y814 into phenylalanine (F814) in a mutant mouse model shows accelerated regenerative responses in an acute skin wound model and post-traumatic OA model (24). However, the cellular mechanism of how mutation of gp130-Y814/SFK signaling diminishes inflammatory responses was unknown. In the present study, we have found that F814 mutant mice are protected from systemic inflammation induced by a High Fat Diet (HFD), with better outcomes seen in major metabolic organs such as the liver as well as musculoskeletal tissues. We have also discovered that gp130 receptor endocytosis is blunted in F814 mouse cells, but the downstream effects are cell-type-specific. While gp130-Y814 signaling appears to be dispensable in monocytes/macrophages, F814 mutation and subsequent blunted gp130 endocytosis prevent chondrocytes from activating the catabolic p38 pathway and secreting cartilage-degrading MMPs, nominating the gp130/SFK/endosomal signaling axis as a potentially novel therapeutic target for chronic inflammatory diseases.

## Results

### Genetic mutation of gp130-Y814 ameliorates systemic inflammation induced by HFD

Based on our previous data showing that F814 mice have enhanced tissue regeneration in a relatively acute inflammatory setting (24), we sought to assess whether these mutant mice are also protected from chronic systemic inflammation using a long-term HFD-induced obesity model (Fig. 1A). WT and F814 mutant mice were given a HFD (60% fat) for 42 weeks, but surprisingly there was no significant difference in total weight gain between the two genotypes (Fig. 1B). DEXA scanning supported this by showing that distribution of the adipose tissue and overall fat content was similar between WT and F814 mice (Fig. 1C). However, when assessing systemic inflammatory factors in the blood serum, circulating CRP and MCP1 were significantly lower in HFD-fed F814 mice (Fig. 1D). While circulating levels of IL-6 were elevated compared to mice fed normal diet (ND), there was not a significant difference between WT and F814 mice fed HFD. Furthermore, when assessing circulating leukocytes in the peripheral blood by white blood cell differential, neither lymphocytes (T and B cells) nor other myeloid cells such as neutrophils were significantly different (Fig. 1E). Again, we detected significantly fewer monocytes in circulation from F814 on a HFD compared to WT (Fig. 1E), suggesting that the monocytes/macrophages from F814 mice may regulate responses to inflammation.

**Figure 1.**
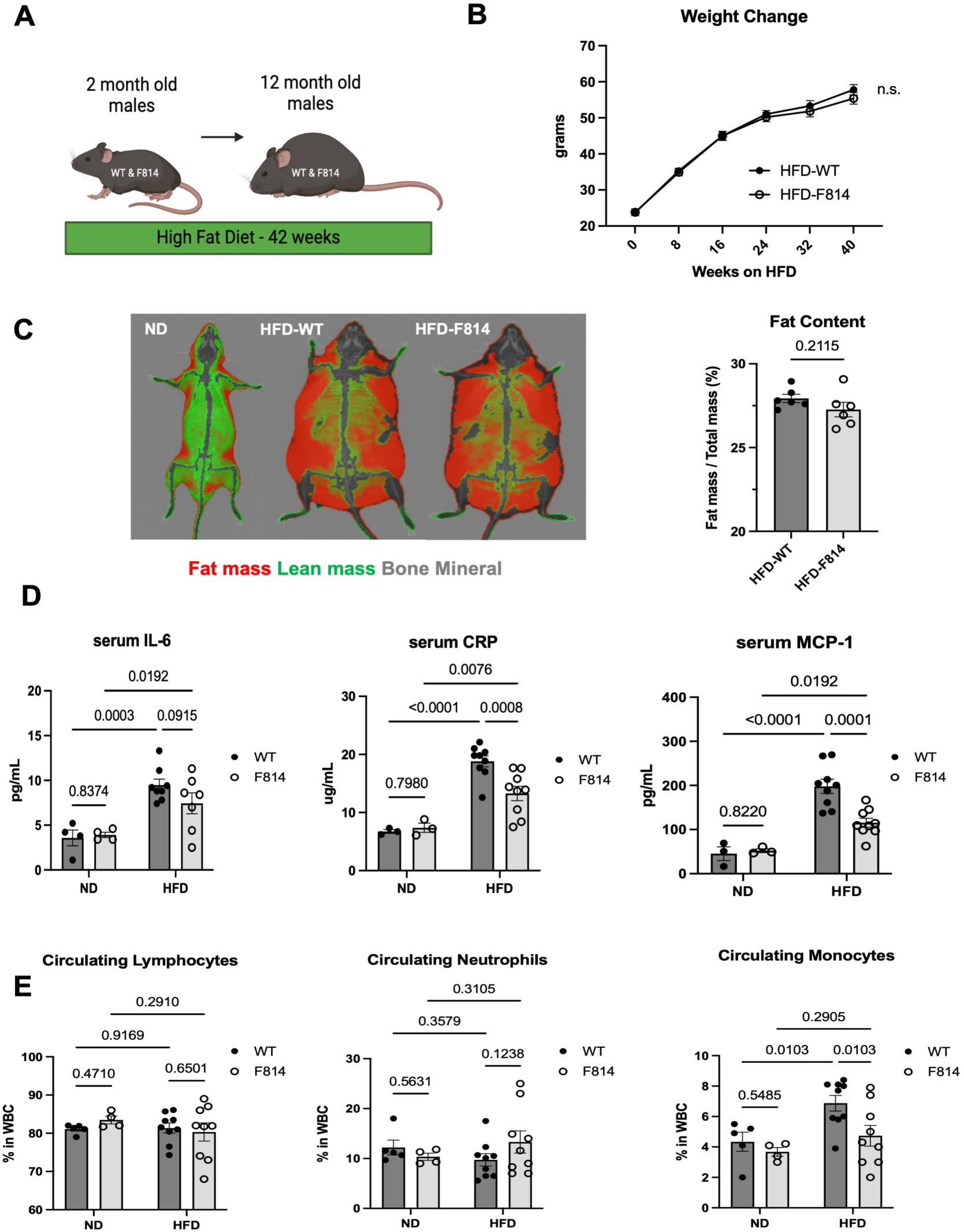
Gp130-Y814 mutant mice have reduced systemic inflammation in response to long-term high fat diet (HFD). (A) Schematic of experimental design. (B) Weight change over time of mice on high fat diet (HFD), n=10 biological replicates per group. (C) Representative images and body composition analysis of WT mice on normal diet (ND) or HFD, generated by DEXA scan. (D) Serum levels of IL-6, CRP, and MCP-1 measured by ELISA in mice fed ND and HFD for 42 weeks. (E) Circulating lymphocytes, neutrophils, and monocytes in peripheral blood from mice on HFD as detected by white blood cell differential. Data points represent biological replicates, error bars are shown as mean +/- SEM, and statistical analyses were performed with 2-way ANOVA with Fisher’s LSD test for more than two groups or Student’s t-test for comparing two groups; p-values less than 0.05 were considered significant.

### F814 mice on HFD show less liver inflammation, fibrosis and myeloid cell infiltration

Given that the liver is a major metabolic organ affected by HFD–induced chronic systemic inflammation, we performed bulk RNA sequencing on liver tissue and evaluated Gene Ontology (GO) terms associated with genes upregulated in HFD-fed WT mice compared to F814 mice (Fig. 2A). Enriched GO terms such as tissue remodeling, endocytosis, and regulation of myeloid leukocyte differentiation were upregulated in HFD-fed WT mouse livers relative to HFD-fed F814 mouse livers. Notably, genes associated with inflammation (*Ccl28, Ccl6, Tlr2*) and tissue remodeling (*Col1a1, Mmp2, Spp1*) were upregulated in the HFD-fed WT compared to HFD-fed F814 mouse livers (Fig. 2B). We histologically confirmed evidence of fibrotic tissue remodeling in the livers via Picrosirius Red staining (Fig. 2C) and monocyte/macrophage presence via CD68 staining (Fig. 2D), whereby F814 mouse livers exhibited markedly less fibrosis and monocyte/macrophage presence compared to WT mouse livers in response to HFD.

**Figure 2.**
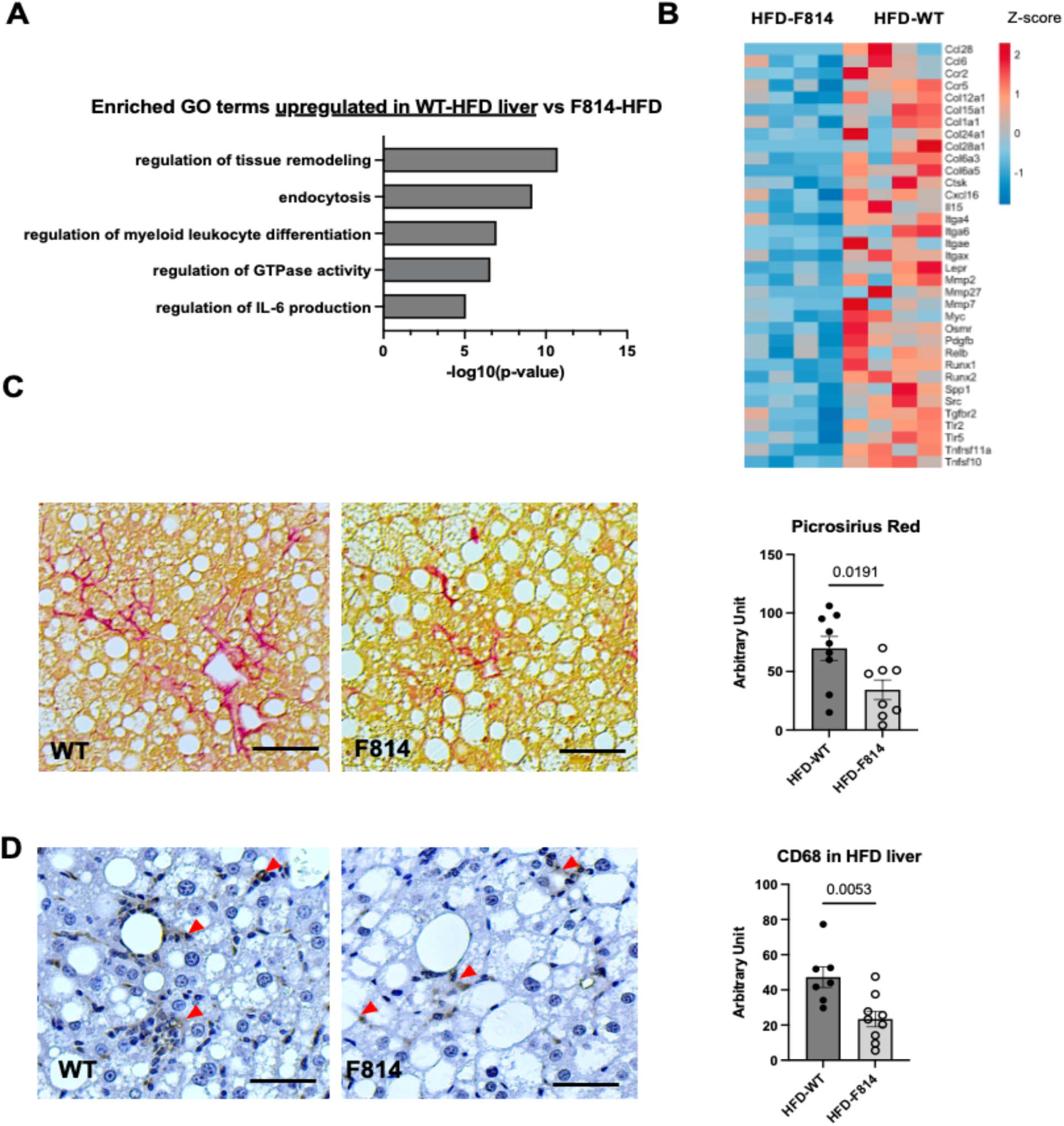
Livers from F814 mice show decreased fibrosis and inflammatory response to HFD compared to WT mice. (A) Enriched Gene Ontology (GO) terms from bulk RNA sequencing of WT HFD livers compared to F814 HFD livers; log2 FC>0.5 and p-adj.<0.01 were considered significant. (B) Heatmap of differentially expressed genes comparing HFD livers of F814 versus WT mice, n=4 biological replicates. (C) Representative images of livers from WT and F814 mice on HFD stained with Picrosirius Red to indicate fibrosis (red staining). (D) Immunohistochemical staining of HFD livers for macrophages (CD68; red arrows). Scale bars represent 20 um. (E) Quantification of (C) and (D); data points represent biological replicates, error bars are shown as mean +/- SEM. Statistical analyses were performed with Student’s t-test; p-values less than 0.05 were considered significant.

### Adipose tissue from HFD have fewer myeloid cells in F814 mice

Another major metabolic organ responsive to HFD is adipose tissue (25). Therefore, we assessed the adipose tissue from mice fed the HFD and found that while *IL-6* transcript levels were not significantly different between WT and F814 mice, classical pro-inflammatory gene *Tnfα* was lower in F814 mice (Fig. 3A). Other genes such as *Ccl2* (MCP-1) and *Spp1* were also upregulated in WT mouse adipose tissue compared to F814 mouse adipose tissue (Fig. 3A). Similar to the liver, we also detected fewer monocytes/macrophages within F814 adipose tissue via CD68 staining (Fig. 3B), which was validated by flow cytometry to show less Ly6C^+^ monocytes in F814 adipose tissue from HFD-fed mice (Fig. 3C).

**Figure 3.**
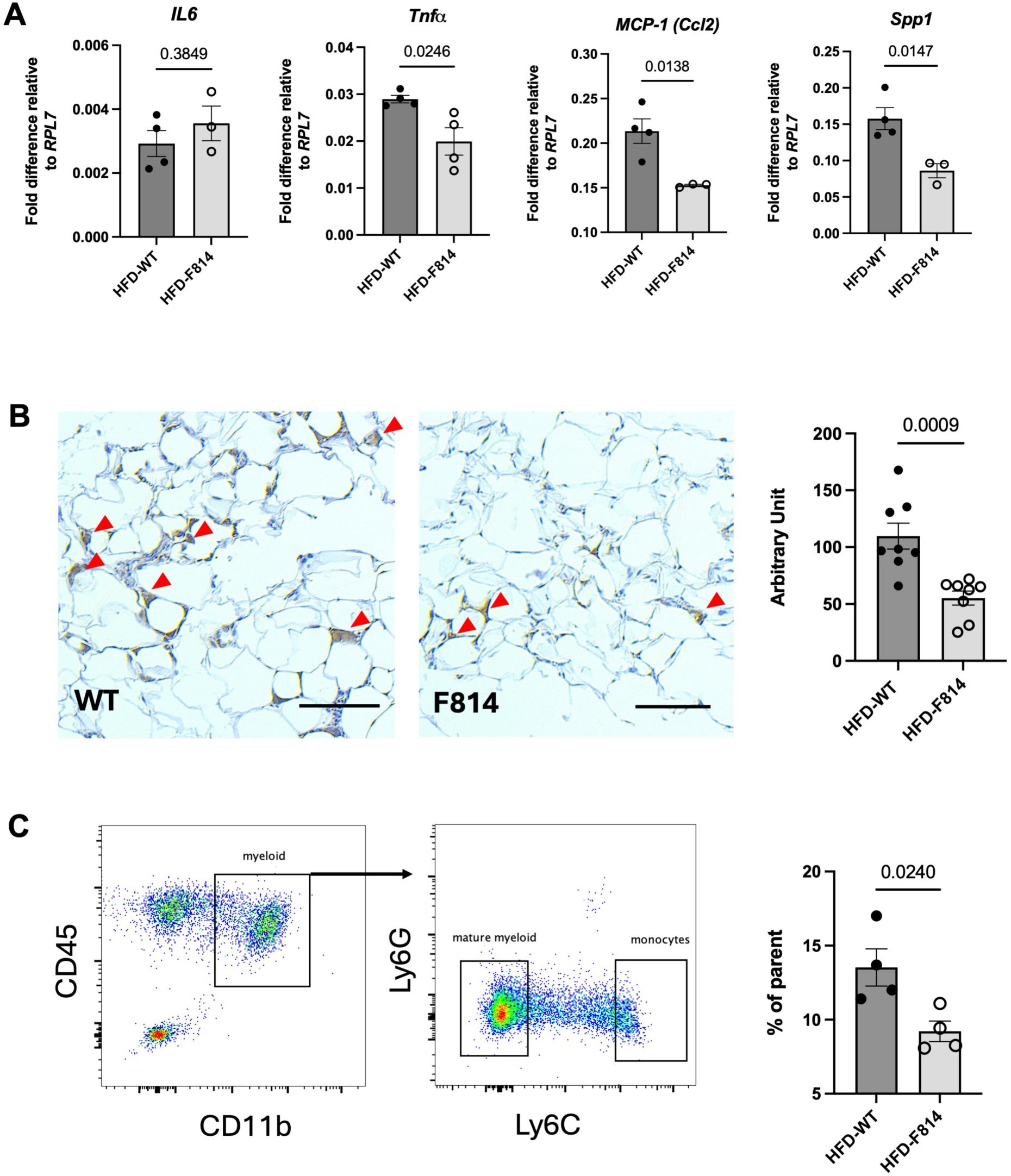
Adipose tissue and peripheral blood from F814 on HFD show less monocytes compared to WT. (A) Transcript levels of genes in adipose tissue of HFD mice measured by qPCR. (B) Representative images and quantification of adipose tissue from HFD mice stained for macrophages (CD68, brown; red arrows). Scale bars represent 10 um. (C) Flow Cytometric analyses of monocytes detected in HFD adipose tissue. Data points represent biological replicates, error bars are shown as mean +/- SEM, and statistical analyses were performed with 2-way ANOVA with Fisher’s LSD test for more than two groups or Student’s t-test for comparing two groups; p-values less than 0.05 were considered significant.

### F814 monocytes/macrophages have blunted gp130 endocytosis

Our previous data have shown that reduced activation of pro-inflammatory macrophages monocytes/macrophages is one of the most prominent phenotypic findings in F814 mice (24). In the present study, we also observed increased monocyte/macrophage presence in both circulation and in major metabolic organs of WT mice compared to F814 mice on a HFD. Therefore, we predicted that the Y814 mutation may bias the functional trajectory of monocytes/macrophages. As bone marrow-derived monocytes become pro-inflammatory monocytes/macrophages in response to inflammation (26), we began by sorting Ly6C^+^ bone marrow monocytes from WT and F814 mice, where monocytes could be separated by surface gp130 levels (Fig. 4A, left). Gp130 expression correlated with Ly6C^+^ levels, suggesting that gp130 is enriched in a pro-inflmmatory subtype of monocytes (27) (Fig. S1A). We then performed bulk RNA sequencing to compare upregulated GO terms between monocytes that express high levels of gp130 versus gp130-low monocytes within each genotype (Fig. 4B, Fig. S1B). Upregulated GO terms in WT gp130-high monocytes included leukocyte activation, inflammatory responses and unexpectedly, endocytosis compared to their gp130-low counterpart. In contrast, F814 gp130-high monocytes had upregulated terms like cell division and regulation of microtubule-based movement (Fig. 4B). Comparing WT versus F814 gp130-high mouse monocytes yielded fewer significant GO terms (Fig. S1C).

**Figure 4.**
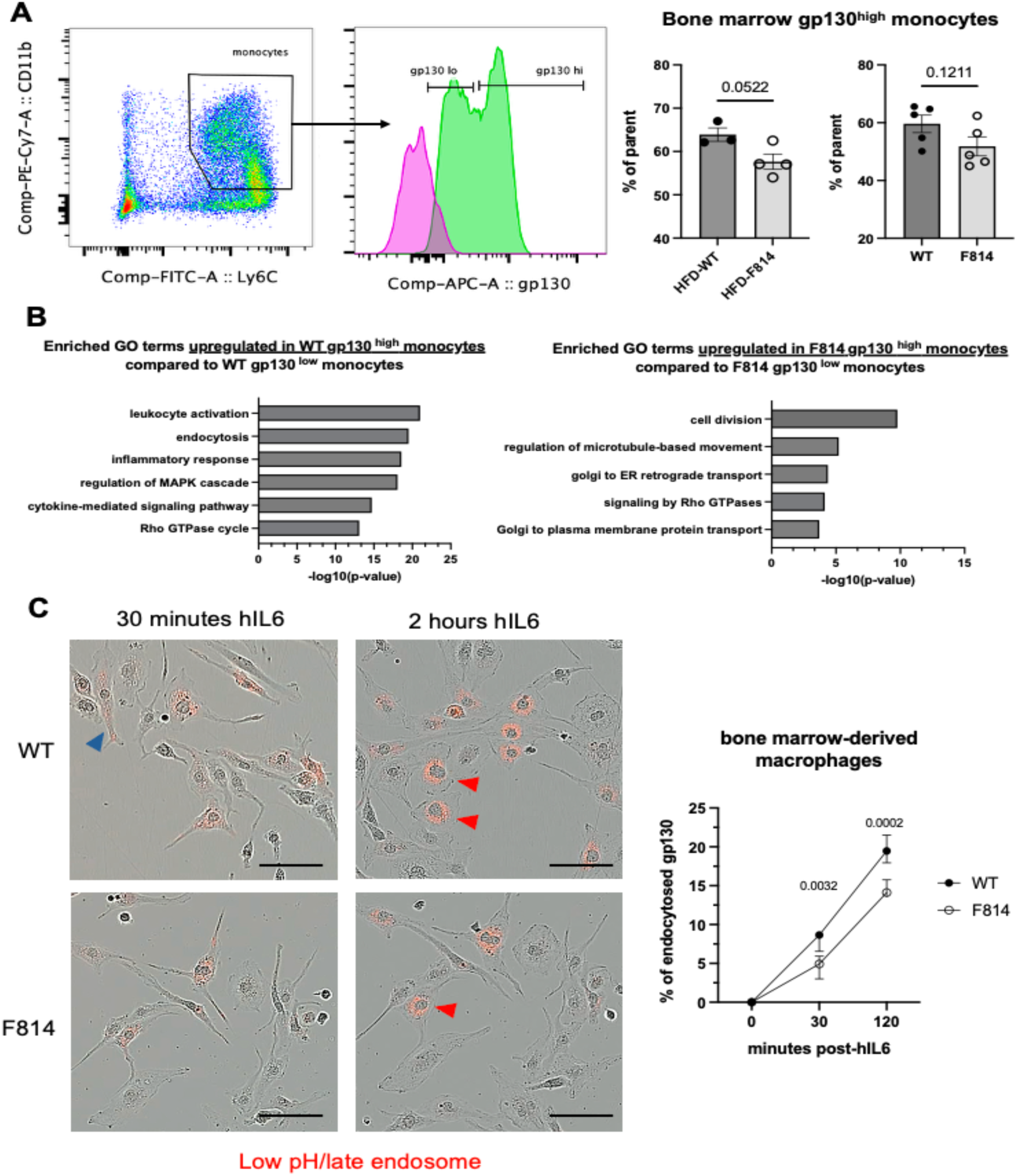
F814 monocytes/macrophages have impaired gp130 endocytosis compared to WT. (A) Flow Cytometric analysis, gating strategy, and quantification of gp130 high and low bone marrow monocytes between F814 and WT mice. Data points represent biological replicates, error bars are shown as mean +/-SEM. Statistical analyses were performed with Student’s t-test; p-values less than 0.05 were considered significant. (B) Enriched Gene Ontology (GO) terms from bulk RNA sequencing of gp130 high monocytes compared to gp130 low monocytes between WT and F814 bone marrow monocytes; log2 FC>0.5 and p-adj.<0.01 were considered significant. (C) Representative images of gp130 internalization in bone marrow-derived macrophages and quantification using Incucyte; red arrows indicate late endosomal/lysosomal gp130. Scale bar represents 20 um, n=3 biological replicates, error bars are shown as mean +/- SEM and statistical analyses were performed with 2-way ANOVA with Fisher’s LSD test.

We tested the two most upregulated GO terms (leukocyte activation and cell division) in gp130-high monocytes from WT and F814 mice;however, when assessing classical monocyte/macrophage pro- and anti-inflammatory gene activation in response to IL-6, there were no significant differences except for *Nos2* (iNOS), which we have previously reported (24) (Fig. S2A). When assessing cell division via BrdU proliferation assay, F814 gp130^+^ monocytes were not significantly more proliferative than their WT counterparts (Fig. S2B). Interestingly, the second most upregulated GO term in WT gp130-high monocytes was endocytosis, which we also observed in HFD-fed WT mouse livers compared to HFD-fed F814 mouse livers (Fig. 2A).

Previous groups have shown that cell surface levels of IL-6 receptor, IL-6Rα, are relatively low in most cell types (28), so we used a soluble IL-6 + IL-6Rα fusion protein, termed hyper IL-6, to stimulate the myeloid cells (29). When we tested gp130 endocytosis on monocytes/macrophages via flow cytometry after hyper IL-6 stimulation, we indeed observed significantly fewer surface level gp130 on WT mouse cells compared to F814 at 30 minutes (Fig. S2C). We further validated these results using Incucyte^TM^, where we were able to observe intracellular spatial localization of a pH-sensitive-labeled gp130 antibody over time (Fig. 4C). At 30 minutes post-hyper-IL6 treatment, we observed increased gp130 localization in early endosomes of WT mouse cells. This localization was sustained at 2 hours, with significantly more WT cells exhibiting gp130 in late endosomes/lysosomes compared to F814 cells (Fig. 4C). This suggested that the rate of gp130 endocytosis may be a cellular mechanism that influences the functional output of monocytes/macrophages.

Because F814 mice carry the Y814 mutation in all cell types, we sought to determine whether the mutation of Y814 specifically in monocytes and macrophages is sufficient to alter their functional phenotype, independent of contributions from mesenchymal cells, which also harbor the F814 mutation. We used a bone marrow transplantation model with a genetic reporter (LysM-Cre^ERT2^/tdTomato/gp130^WT/WT^ or LysM-Cre^ERT2^/tdTomato/gp130^F814/F814^) to assess how restriction of F814 to only hematopoieitc cells influences their ability to become pro-inflammatory (Fig. 5A). Donor WT or F814 mouse bone marrow cells (CD45.2^+^) were transplanted into CD45.1^+^ recipients, and after the recipients recovered, we subjected them to an acute model of arthritis induced by local injection of the Complete Freund Adjuvant (CFA) (30). Two days post intra-articular joint injection, we sorted tdTomato^+^ donor monocytes/macrophages that migrated into the joint (Fig. 5B) and assessed their gene expression signature. However, to our surprise, there were no significant differences in pro-inflammatory genes between transplanted WT and F814 monocytes/macrophages (Fig. 5C). This suggested that F814 mutation/gp130 endocytosis in monocytes/macrophages was not a key driver of favorable phenotypes seen in the global F814 mice. Instead, F814 mutation and gp130 endocytosis in other cell types, such as mesenchymal cells, may play a more significant role in driving these outcomes and indirectly regulating the activation of innate immune cells.

**Figure 5.**
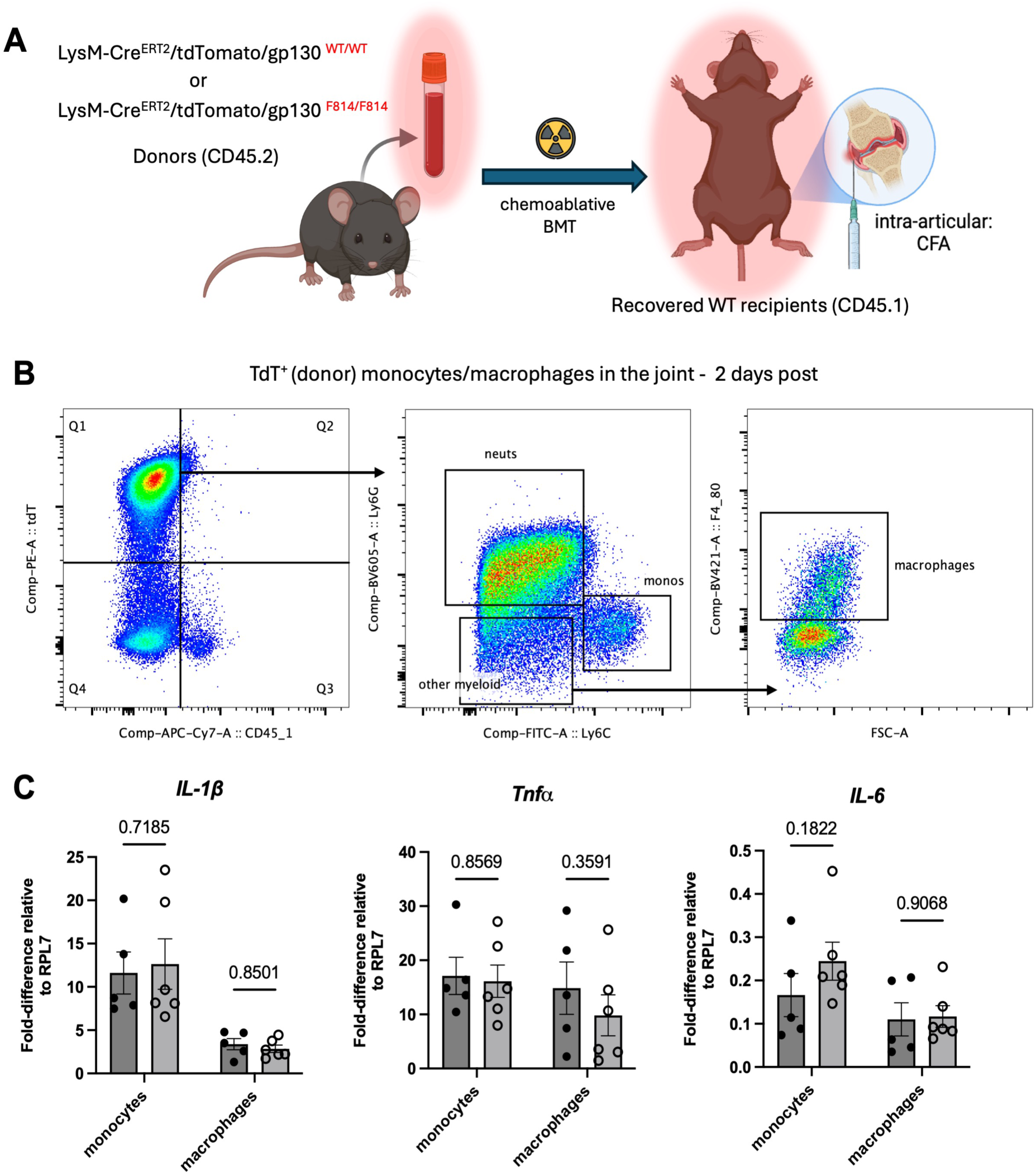
F814 mutation in hematopoietic cells does not influence monocyte/macrophage functional phenotype. A) Schematic of experimental design for genotypes, bone marrow transplantation (BMT), and acute inflammatory arthritis modeling. B) Representative FACS gating strategy for donor-derived tdTomato+ monocytes/macrophages from the joint 2 days post CFA injection. C) Transcript levels of traditional pro-inflammatory genes in sorted donor-derived monocytes/macrophages measured by qPCR. Data points represent biological replicates, error bars are shown as mean +/- SEM, and statistical analyses were performed with 2-way ANOVA with Fisher’s LSD test; p-values less than 0.05 were considered significant.

### F814 mice show resistance to bone and cartilage loss induced by HFD

We then switched focus to musculoskeletal-specific cells, given that the link between chronic systemic inflammation and obesity-induced arthritis has been described . When we assessed overall bone density in the long bones of WT and F814 mice on a HFD, we saw significantly more trabecular bone density in F814 mice (Fig. S3A). Given that bone density is regulated by the balance of osteoclasts (specialized monocyte-derived bone resorbing cells) and osteoblasts (mesenchymal stromal cell-derived bone secreting cells) (31–34), we assessed the ability of WT and F814 mouse cells to form osteoclasts. We did not detect any significant morphological (Fig. S3B) or hallmark osteoclast gene differences (Fig. S3C) between the two genotypes, supporting our findings that F814 mutation has little effect on the functional phenotype of osteoclasts. We did, however, see less Tartrate-Resistant Acid Phosphatase (TRAP) staining, a marker for osteoclast activity, in the long bones of HFD-fed F814 mice (Fig. S3D). gp130 signaling in bone marrow stromal cells, osteoblasts, and osteocytes induces RANKL expression, thereby promoting osteoclastogenesis (35,36). When we assessed gene levels of RANKL, the ligand necessary for osteoclast formation and mainly produced by mesenchymal stromal cells (37), HFD-fed F814 mouse cells had significantly lower expression of the *RANKL* gene compared to WT (Fig. S3E).

Furthermore, when observing the gross morphology of the joints after HFD, we noticed significantly more preserved articular cartilage and better OARSI arthritis scoring in F814 mice (Fig. 6A), suggesting that F814 signaling may be more essential in chondrocytes and has mitigated HFD-induced osteoarthritis (OA).

**Figure 6.**
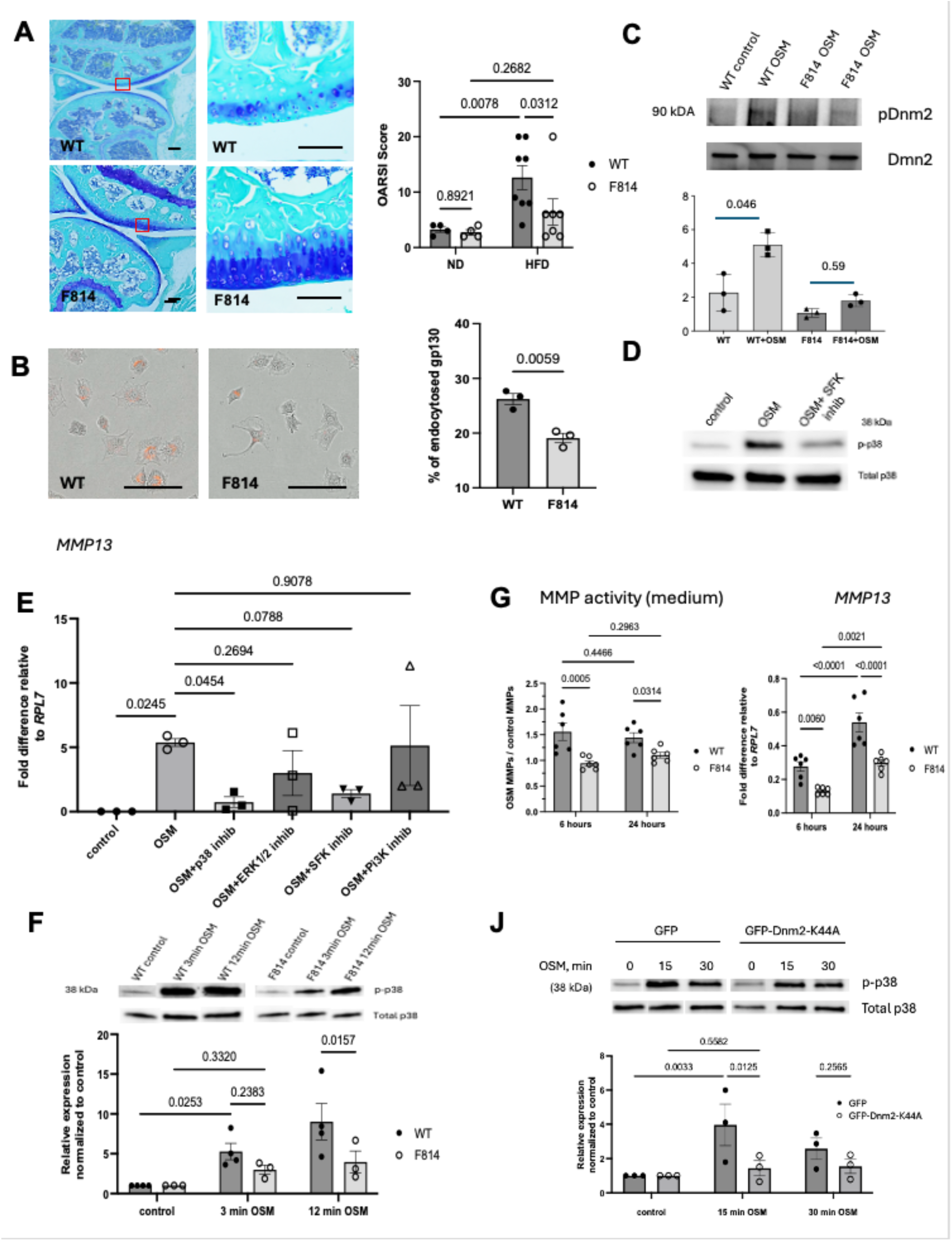
F814 mice are protected from HFD-induced degenerative joint disease and show reduced cellular stress response to IL-6 cytokine OSM. (A) Representative images of joint articular cartilage via Toluidine Blue staining to assess glycosaminoglycan levels (blue); Osteoarthritis scoring via OARSI standards defined in the Methods section. Red cutout shows higher magnification and scale bar represents 20 um. (B) Representative images of gp130 internalization in chondrocytes after OSM stimulation and quantification (bottom) using Incucyte, scale bar represents 20um. (C) F814 chondrocytes show reduced activation of GTPase Dynamin 2 by OSM. (D) Inhibiting the SFK pathway or endocytosis blunts downstream p38 activation in chondrocytes. Representative Western blot of phospho- and total p38 after SFK inhibitor and OSM stimulation for 15 minutes in pig chondrocytes. (E) Transcript levels of *MMP13* in pig chondrocytes with various downstream pathway inhibitors after OSM stimulation for 24 hours as measured by qPCR. (F) MMP release assay detecting MMPs activity reflecting the protease levels secreted by chondrocytes into cell culture media after OSM stimulation. Transcript levels of *MMP13* in WT and F814 chondrocytes measured by qPCR. (G) Representative Western blot of phospho- and total MAPK/p38 in WT versus F814 chondrocytes and quantification (bottom). (J) Representative Western blot of phospho-and total MAPK/p38 in chondrocytes trasduced by control (GFP) or Dnm44A after stimulation with OS. Quantification is presented at the bottom. Data points represent biological replicates, error bars are shown as mean +/- SEM, and statistical analyses were performed with 2-way ANOVA with Fisher’s LSD test; p-values less than 0.05 were considered significant.

The levels of inflammation in the synovioum was low in both genotypes as determined by the histological assessment suggesting a degenerative rather then an inflammatorynature of the observed OA (Fig.S4). Because of our advanced-level expertise in this field, we next focused on a more detailed characterization of the signaling mechanisms leading to reduced cartilage loss in F814 mice. To validate that blunted gp130 endocytosis was a universal cellular mechanism in F814 mice, we measured gp130 internalization in chondrocytes (Fig. 6B) and saw significantly less endocytosed gp130 in F814 compared to WT chondrocytes. Further validation with immunocytochemistry showed less co-localization of gp130 with LAMP1, a late endosome/lysosome marker, in F814 chondrocytes stimulated with OSM (Fig. S5). Comparative studies by our group has previously shown OSM to be the most catabolic IL-6 cytokine for chondrocytes inducing massive MMP13 expression and matrix breakdown. Our group has previously shown that OSM-induced gp130-Y814 signaling activates SFKs via phosphorylation and that F814 mutation blunts SFK activation (24). Additionally, other studies have demonstrated that SFK activity is required to activate GTPases such as dynamins (*Dnm*), which are essential for receptor-mediated endocytosis. Mammalian genome contains three *Dnm* genes (38). *Dnm1* is selectively expressed at high levels in neurons, *Dnm2* is expressed ubiquitously, and *Dnm3* is found predominantly in the brain (at much lower levels than *Dnm1*) and testis (39,40). Accordingly, our studies have focused on *Dnm2* (encoded by the *Dnm2* gene). Dnm2, a 90-100 kDa GTPase, plays a role in receptor-mediated endocytosis via membrane fission and is a direct target of SRC (41). SFK and Dnm2 unit is also required for the packaging of cargo-loaded exosomes in the Golgi apparatus (42,43), and this is a critical step required for the secretion of biologically active stress-response-related molecules, such as proteases, by stress-activated cells.

We next assessed levels of active Dynamin 2 (Dnm2) in chondrocytes stimulated with OSM to link gp130 signaling, SFK activation, and endocytosis as a proposed downstream mechanism to explain the phenotype observed in F814 mice. Dnm2 is a ubiquitously expressed isoform and it has previously been shown to be activated by SFK in various cell types (42,44). We found that stimulated F814 mouse chondrocytes had significantly less active Dnm2 compared to WT chondrocytes (Fig. 6C), suggesting Dnm2 activation as a downstream mechanism of gp130/SFK signaling and that activation of this mechanism by IL-6 cytokines is markedly reduced in F814 mutant cells resulting in significantly decreased gp130 internalization.

Our molecular data demonstrated that pharmacological inhibition of SFKs leads to: (a) a significant reduction in MAPK/p38 activation in response to OSM (Fig. 6D), and (b) that pharmacological inhibition of MAPK/p38 yields the most pronounced protective effect against OSM-induced secretion of the catabolic enzyme MMP13 in chondrocytes. (Fig. 6E). MMP13 is a well-known direct target of p38/MAPK (45,46). Moreover, MAPK/p38 activity in chondrocytes has already been linked to gp130 endocytosis (47), and activation of catabolic and stress pathways, including MMP production (48). Based on these observations, we hypothesized that reduced internalization of gp130 in the F814 mutant will lead to reduced MAPK/p38 activation in response to OSM. To test that, we assessed pMAPK/p38 activity in F814 mouse chondrocytes stimulated with OSM and observed a marked suppression of phosphorylated MAPK/p38 in the treated F814 mouse chondrocytes compared to the WT (Fig. 6F). This decrease correlated with reduced enzymatic activity of MMPs in the culture media after OSM treatment (Fig. 6G-left), which was consistent with the observed inhibition of Dnm2 activity. SFK and Dnm2 unit is also required for the packaging of cargo-loaded exosomes in the Golgi apparatus (49,50) and this is a critical step required for the secretion of biologically active stress-response-related molecules, such as proteases, by stress-activated cells. *The Mmp13* gene expression levels from the same cells supported these results (Fig. 6G-right), highlighting how blunted gp130 endocytosis in F814 chondrocytes can affect downstream pathways and functional outcomes.

Finally we tested whether inactivation of the endogenous Dnm2 activity by overexpression of a dominant-negative Dnm2 (Dnm2-K44A) can reduce the activation of p38 kinase by OSM. This approach has previously been shown to mitigate activation of MAPK/p38 kinase by fibronectin fragments in chondrocytes and reduce secretion of catabolic proteases such as MMP13 (51). In good agreement with these studies, we have shown that Dnm2-K44A has significantly reduced MAPK/p38 activation by OSM nominating Dnm2 as one of the key regulators of pro-inflammatory and pro-catabolic signaling by various upstream mediators, including the IL-6 family of cytokines (Fig. 6J). Based on these findings, Dnm2 may represent an attractive therapeutic target for chronic inflammatory and degenerative diseases such as osteoarthritis.

## Discussion

We and others have described gp130 signaling as a complex multimodal sensor that finely tunes multiple signaling pathways to be either pro-regenerative or pro-inflammatory (52,53). In this study, we have shown that gp130 F814 mice are protected from chronic systemic inflammation in multiple organs. It was surprising to observe little overall difference in weight gain and fat content between WT and F814 mice, as previous work has described the leptin/gp130 relationship as a potential therapeutic target (54) and that the IL-6 deficient mice are obese (55). Even more interesting were insignificant differences between systemic levels of IL-6, yet markedly less circulating levels of CRP and MCP-1. IL-6 is recognized as an upstream activator of both CRP and MCP-1 production in the liver and other tissues (56,57), therefore, the lower levels of both factors in F814 mouse serum cannot be explained by lower levels of circulating IL-6.

Given that the liver and adipose tissue are the two major metabolic organs that respond to HFD and chronic systemic inflammation (58), F814 metabolic tissues had fewer detected adverse morphological and genetic profiles. It was interesting that GO terms such as “endocytosis”, “regulation of myeloid leukocyte differentiation”, and “regulation of GTPase activity” were upregulated in HFD-fed WT livers compared to F814. This aligns with our previous observations of reduced pro-inflammatory myeloid cells in single-cell RNA sequencing of acute skin wounds in F814 (24). Bulk RNA sequencing was performed on total livers, which include both liver-specific macrophages (Kupffer cells), hepatocytes, and surrounding mesenchymal stromal cells (59), so it is difficult to claim which cell type contributed the most to “endocytosis” and “regulation of GTPase activity” GO terms. As increased MCP-1 in the adipose tissue has been considered pro-inflammatory and contributes to insulin resistance (60), significantly fewer *MCP-1* transcript detected in F814 on a HFD was an indication that gp130-Y814 mutation can modulate MCP-1 production, which could explain the markedly less monocyte/macrophage populations infiltrating into the tissue. Less systemic MCP-1 detected in the serum could also explain less circulating monocytes detected in F814 on a HFD compared to WT. We were not surprised to see insignificant change in circulating lymphocytes (B and T cells) and myeloid neutrophils between the two genotypes as our previous datasets showed little change in lymphocytes that respond to inflammatory conditions (24) and that mature neutrophils do not express gp130 (61).

Similarly to HFD-fed WT mouse livers, it was interesting to see the “endocytosis” GO term appear in WT gp130-high monocytes; this similarity led us to hypothesize that F814 mutation affects gp130 receptor endocytosis in a more ubiquitous, cell-type-independent manner. Contrary to our original hypothesis and previous indications (24), blunted gp130 endocytosis in F814 monocytes/macrophages did not seem to have a significant effect on the functional output of these cells in the tested assays. When F814 mutation is present only in hematopoietic cells using bone marrow transfer and an acute inflammatory arthritis model, there were no significant differences in F814 monocytes and macrophages to become pro-inflammatory compared to WT mouse cells. This suggests that gp130-Y814 signaling is more important for driving functional responses in mesenchymal and/or stromal cells, not hematopoietic cells, which could partially be explained by inherent levels of gp130 expression between the two cell types – mesenchymal cells such as fibroblasts express higher levels of *Il6st* than hematopoietic cells (Human Protein Atlas) (62). The presented data modified our original hypothesis into the concept that gp130-Y814 signaling is far more important in non-hematopoietic cells in driving pro-inflammatory/pro-catabolic responses and the responses of infiltrating monocytes/macrophages are secondary.

In support of our new hypothesis, mesenchymal cells from the bone marrow and chondrocytes with F814 mutation had significantly reduced catabolic responses in both *in vivo* and *in vitro* assays. F814 mice on a HFD had significantly less osteoarthritis and cartilage degradation, and F814 mouse chondrocytes produced fewer MMPs in response to OSM (24,63), linking blunted gp130 endocytosis to a favorable functional outcome. Cartilage degradation from stressed chondrocytes producing MMPs has been linked to activation of the p38/MAPK pathway (64). The observed marked attenuation of MAPK/ p38 activation in F814 chondrocytes may explain the reduced secretion of MMPs. Our previous studies have shown that chondrocytes from F814 mice have significantly less activation of SFK by IL-6 cytokines, especially OSM. Interestingly, SFKs are known to be one of the major regulators of membrane trafficking and cytoskeletal rearrangement via a direct activation of GTPases including Dynamins as well Rho family of GTPases(44,65).

Interestingly, two independent recent reports have shown the anti-catabolic effects of a dynamin inhibitor, dynasore, in chondrocytes and osteoarthritis (*9*), although this molecule is not selective for dynamin. In one of these studies, the effect was linked to the inhibition of intracellular Notch signaling and was more prominent in aged cells (66). The second report showed reduced levels of reactive oxygen species (ROS) in chondrocytes in the presence of dynasore (*6*). However, dynasore has previously been shown to reduce ROS via a dynamin-independent mechanism (67). Dynasore also reduces labile cholesterol in the plasma membrane, and disrupts lipid raft organization, in a dynamin-independent manner (68).

The presented data suggests a connection between gp130/SFK endosomal signaling, Dnm2 and p38/MAPK pathways, where gp130-Y814 mutation plays a role in blunting gp130 endocytosis. More work is needed to fully define how Y814 controls gp130 endocytosis, but we hypothesize that gp130/SFK activation controls receptor internalization likely through secondary mechanisms such as SFK/dynamin activation to pinch off the endosomal vesicle (69) or SFK/clathrin-related protein activation to form the endosome (70). It also remains unclear why endocytosis is required for the amplification of p38 MAPK signaling and what the precise underlying mechanism might be (71). One of the plausible hypotheses for future studies is that this mechanism is pH dependent; a recent report has demonstrated that SFKs are pH-sensitivekinases and that low pH can activate SFK activity by 10-fold (72). SFKs are known to be highly enriched in endosomes, and it is therefore plausible that some SFK-mediated pathways, such as MAPK/p38, may be amplified in the endosomal environment, which is characterized by a low pH. However, further studies are required to dissect these mechanisms in detail. Our previous work has shown that F814 mutation has much less significant effect on STAT3 activation by IL-6 cytokines. Interestingly, in good agreement with that, previous studies demonstrated that STAT3 activation by IL-6 cytokines is not dependent on endocytosis and therefore selective inhibition of gp130 internalization may be a viable strategy for modulation of the signaling pathways downstream of gp130.

This study has shown, for the first time, that the Y814 residue is directly implicated in the dyn2-mediated internalization of gp130 receptor highlighting gp130-Y814–dyn2 signaling as a key driver of pathological outcomes in chronic inflammatory and degenerative diseases in a cell-type–specific manner. SFK-Dyn2 mediated membrane trafficking and cellular stress responses can be engaged by other cytokine-receptor families, and therefore, the inactivation of this entire effector mechanism may be more potent than the blockade of any of the upstream signals. Although Dnm2 is an attractive therapeutic target, the highly pleiotropic functions of this GTPase make drug development challenging and highly context-specific.

## Supplementary Figures

**Figure S1.**
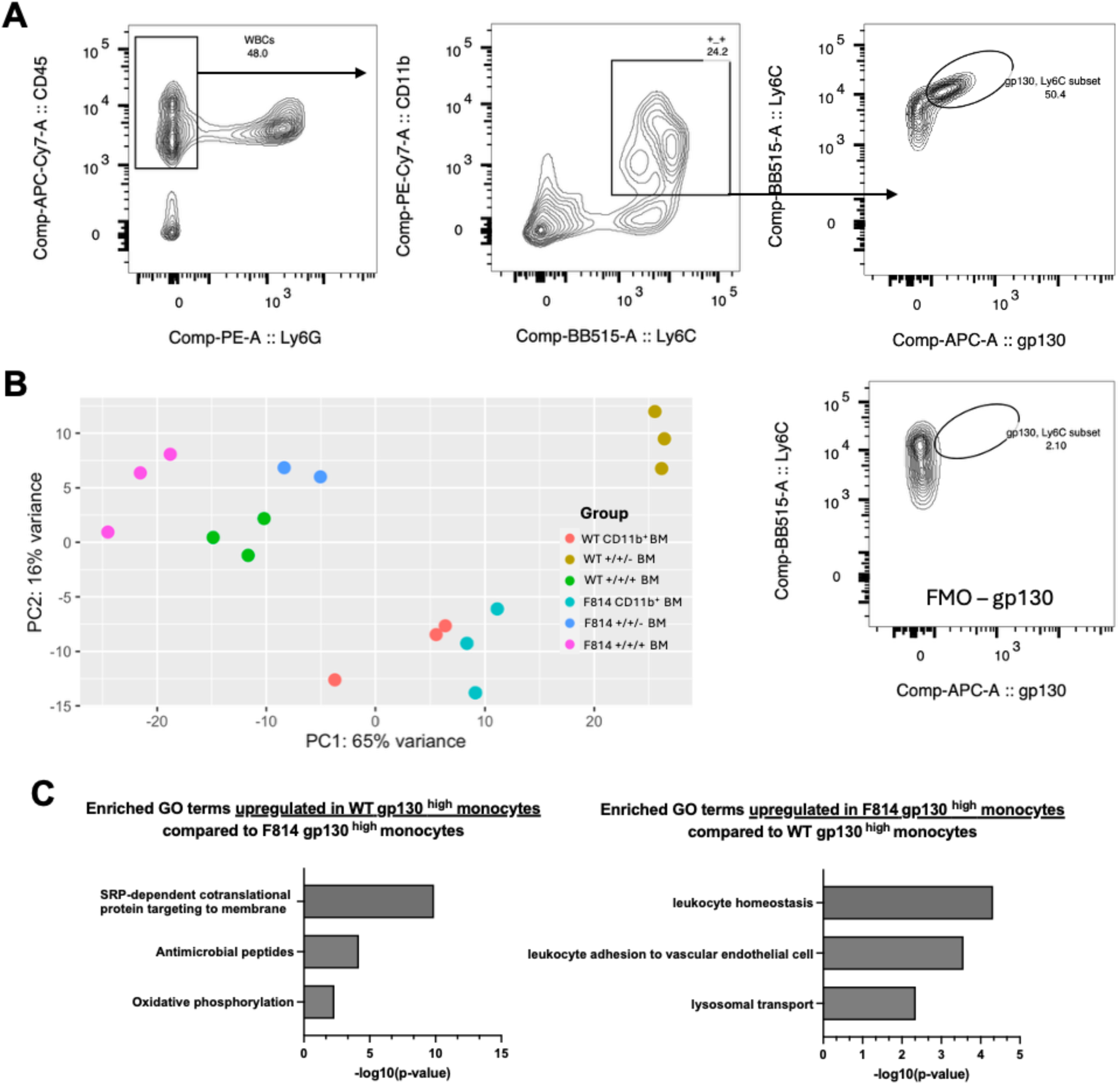
Bulk RNA sequencing of bone marrow gp130 monocytes: gating strategy, PCA analysis, and GO terms. A) Representative flow cytometry gating strategy and Fluorescence Minus One (FMO) controls for sorting gp130+ bone marrow monocytes. B) PCA analysis of bulk RNA-sequencing of bone marrow monocytes. C) Enriched Gene Ontology (GO) terms from bulk RNA sequencing of WT gp130 high monocytes compared to F814 gp130 high monocytes; log2 FC>0.5 and p-adj.<0.01 were considered significant.

**Figure S2.**
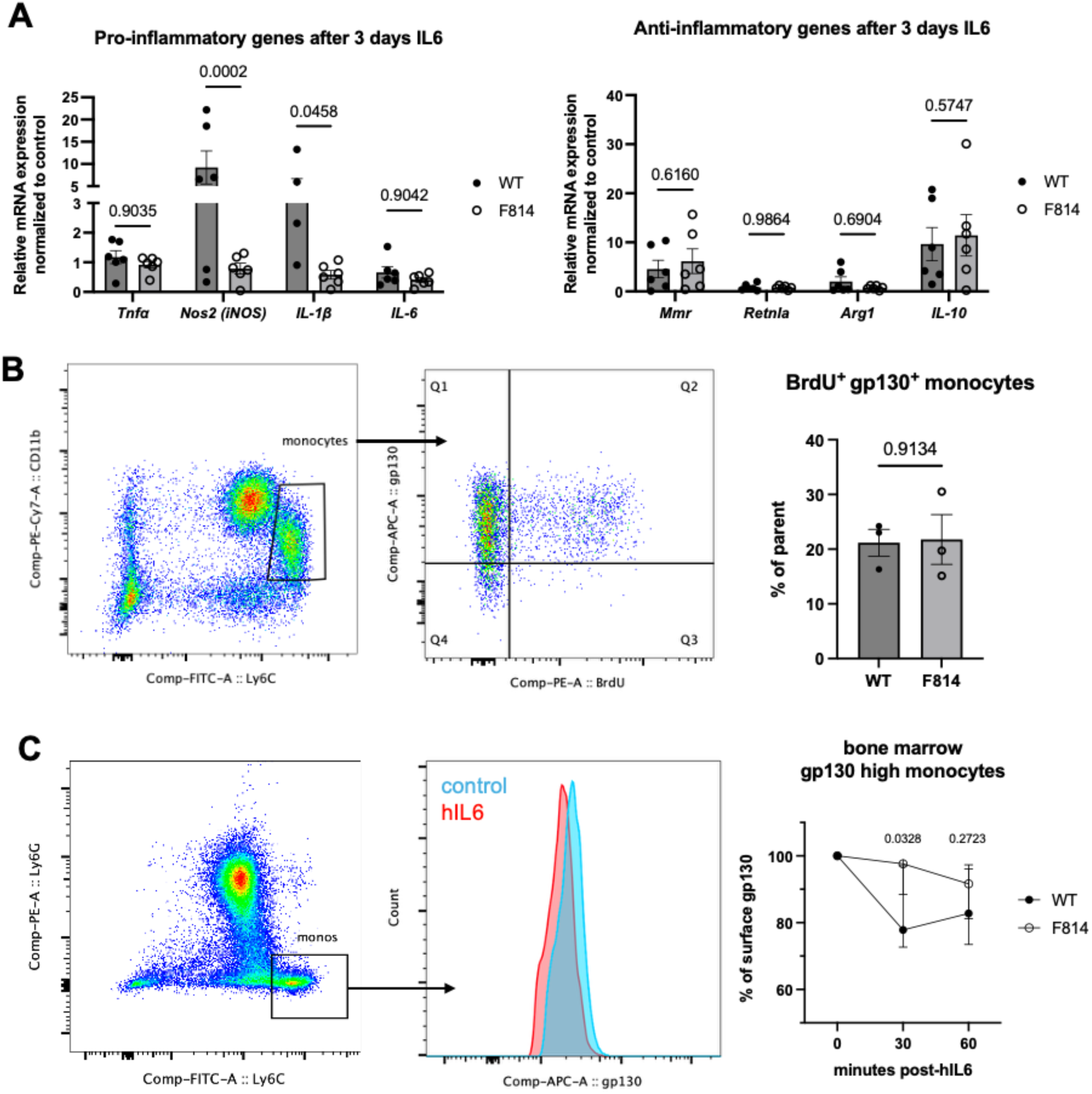
F814 monocytes/macrophages only show modest differences. A) Transcript levels of classical pro-inflammatory and anti-inflammatory genes of bone marrow-derived macrophages after stimulation with IL-6 measured by qPCR. B) Flow cytometric analysis and quantification of BrdU+ (proliferating) F814 and WT gp130+ monocytes. C) Flow cytometric analysis and quantification of gp130 endocytosis on fixed bone marrow cells after stimulation with IL-6 over time; n=3 biological replicates. Data points represent biological replicates, error bars are shown as mean +/- SEM, and statistical analyses were performed with 2-way ANOVA with Fisher’s LSD test for more than two groups or Student’s t-test for comparing two groups; p-values less than 0.05 were considered significant.

**Figure S3.**
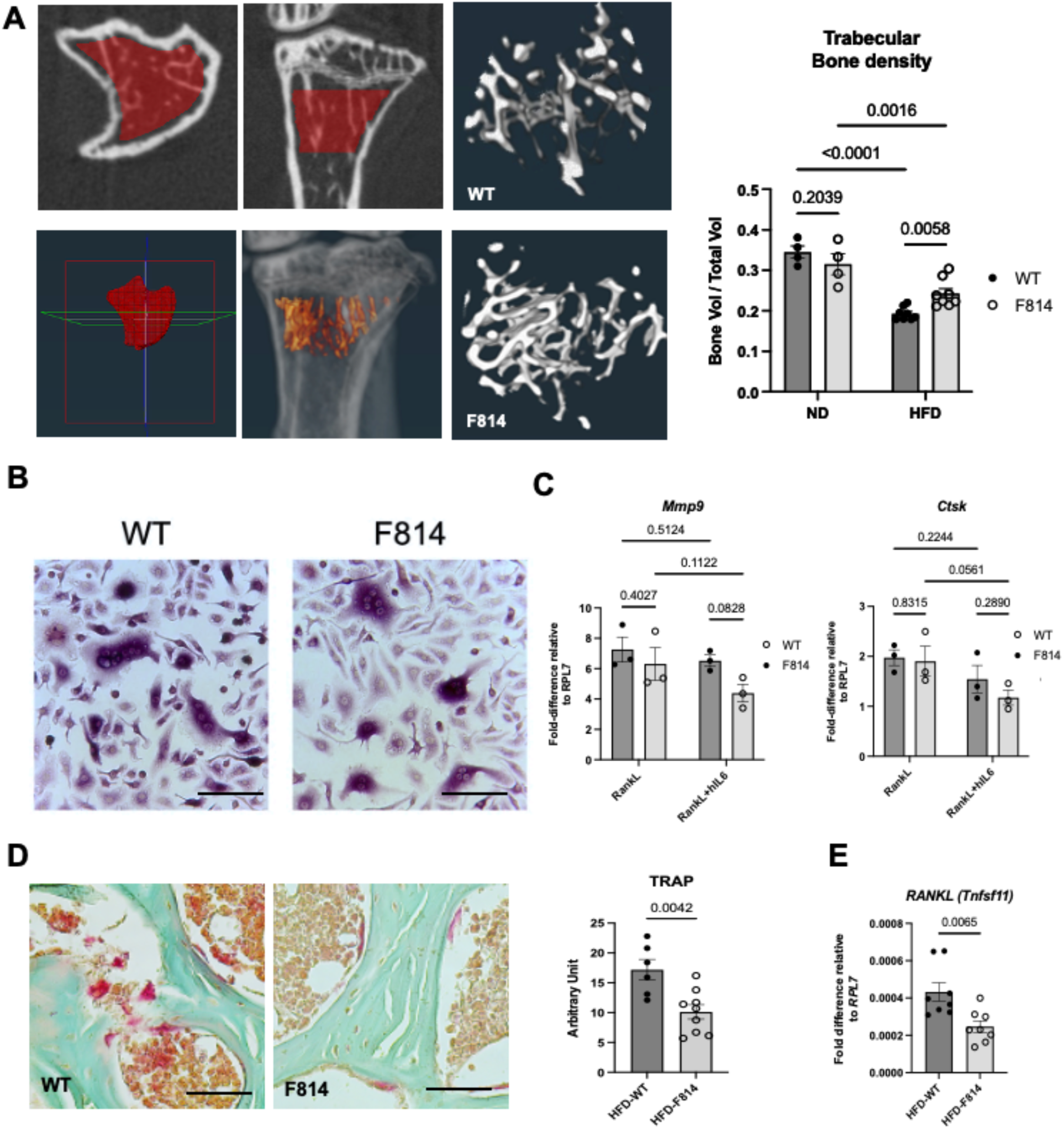
F814 mice on HFD show reduced bone loss compared to WT. (A) Region of interest for micro-CT scans of trabecular bone density and quantitative analysis of WT and F814 mice on HFD.. (B) Representative images of TRAP positive (purple) osteoclasts from WT and F814 mice. (C) Transcript levels of osteoclast marker genes measured by qPCR in osteoclasts. (D) Representative images and quantification of TRAP activity (pink) in trabecular bone (green) of mice on HFD. (E) Transcript levels of osteoclast formation gene *RANKL* in total bone marrow from HFD mice measured by qPCR.

**Figure S4.**
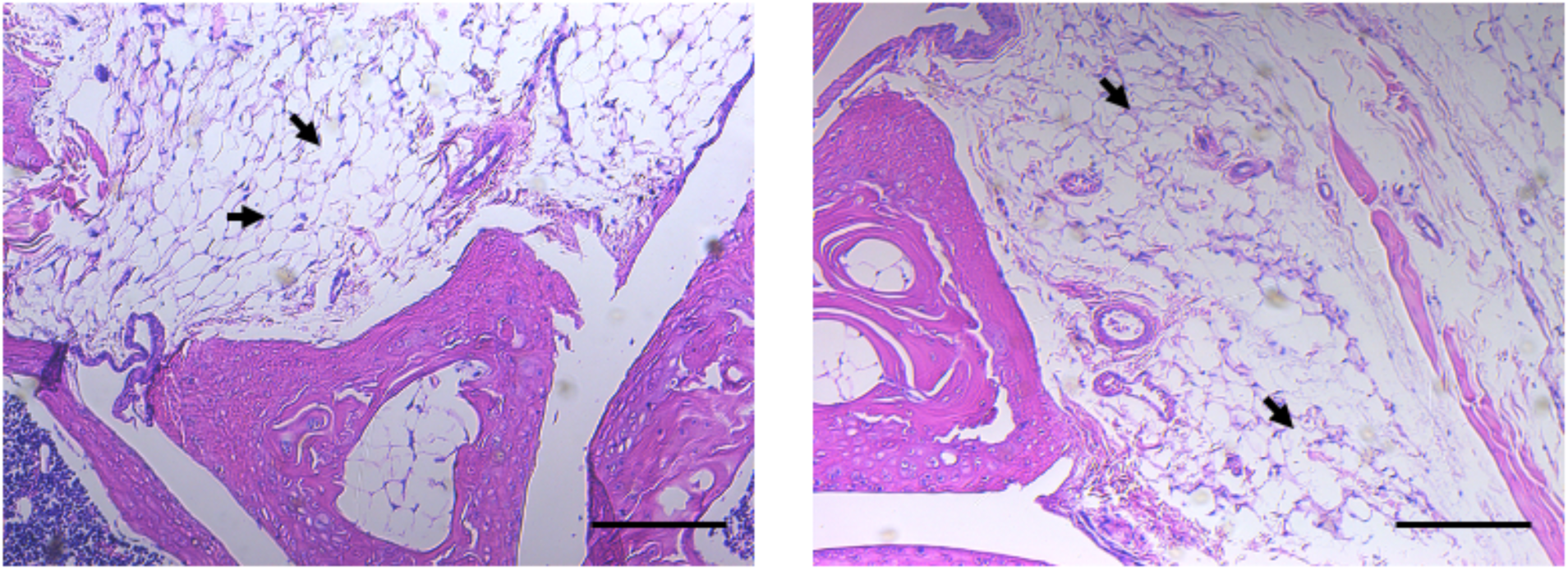
Synovial tissue of the knee joint of experimental mice. Both WT mice and F814 mutants demonstrate minimal levels of synovial inflammation, slight excessive accumulation of adipose tissue is observed (arrows) in both groups exposed to the High Fat Diet. Hematoxylin and eosin (H&E) staining was performed. Representative images are shown. Scale bars=100 μm.

**Figure S5.**
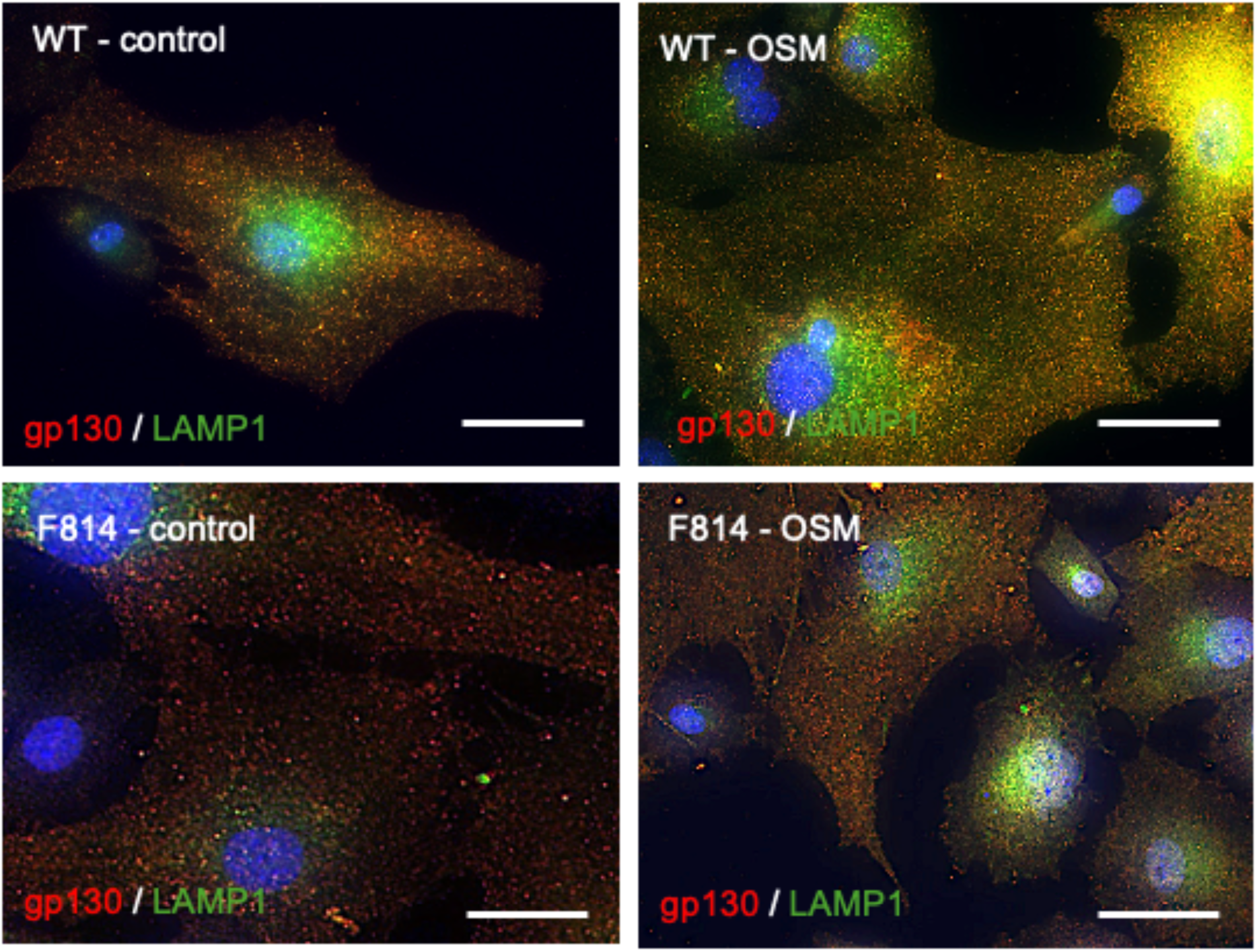
F814 chondrocytes show altered gp130 endocytosis and downstream signaling pathways. Representative images of chondrocytes stained with DAPI (blue), gp130 (red), and LAMP1 antibodies (green) before and after stimulation with OSM.Scale bar =10 μM.

## Methods

### Mouse genetics and in vivo modeling

All experimental procedures were approved by the Institutional Animal Care and Use Committee of the University of Southern California and met or exceeded the requirements of the Public Health Service/National Institutes of Health and the Animal Welfare Act. Homozygous mutant gp130-Y814 mice (F814) on a C57BL/6J background were generated using CRISPR-Cas9, and C57BL/6J mice (JAX #000664) were used as wildtype (WT) controls as previously described (24).

For the long-term high fat diet (HFD) model, F814 mutant and WT mice were fed a 60% high-fat diet (Envigo, TD.06414) from 2-months-old for 10 months. Normal diet control mice were fed a standard rodent diet that contains 13% fat (PicoLab, 5053). F814 and WT mice on a normal diet or HFD were sacrificed at 12-months-old.

For genetic studies, tamoxifen-inducible LysM-Cre^ERT2^ mice (JAX #032291) we crossed to Rosa26-tdTomato reporter mice (JAX #007914) to create LysM-Cre^ERT2^/tdTomato/gp130^WT/WT^ mice. Crossing LysM-Cre^ERT2^/tdTomato/gp130^WT/WT^ mice to homozygous F814 mice (gp130^F814/F814^) for at least two generations created LysM-Cre^ERT2^/tdTomato/gp130^F814/F814^ reporter mice, where these mice express the CD45.2 variant. To activate the tdTomato reporter, mice received Tamoxifen (100 μg per gram of body weight) in corn oil via intraperitoneal injection two consecutive days before *in vivo* modeling, and once weekly for the duration of the experiment. For bone marrow transplantation studies, recipient WT mice with the CD45.1 variant allele was purchased from Jackson Labs (JAX #002014).

For the bone marrow transplantation (BMT) model, we used techniques previously described by the Lu Lab at USC (73). Briefly, hematopoietic stem cells [Lineage (CD3, CD4, CD8, B220, Gr1, Mac1, Ter119)^-^ / cKit^+^ / Sca1^+^ / Flk2^-^ / CD34^-^ / CD150^+^] were isolated via FACS from crushed bones of donor mice after enrichment using MACS with CD117-microbeads (Miltenyi #130-091-224). Recipient mice received an intraperitoneal injection of busulfan (chemoablative drug, Sigma #150606) at a dose of 50 mg per kg body weight 24 hours before transplantation. Donor HSCs were transplanted via retro-orbital injection. Donor chimerism in recipients was evaluated 1 and 2 months post BMT.

After full recovery from BMT (3 months post transplant), mice were subjected to an acute inflammatory intra-articular model, where a single intra-articular injection of Complete Freund’s Adjuvant (CFA) was performed using previously described methods (74). Mice were anesthetized with isoflurane and received 30 μl (1 μg/μl) CFA into the joint cavity using a 30g insulin needle. CFA-injected joints were harvested 2 days post injection.

### Serum ELISAs

Mouse blood was collected from the submandibular vein and left to clot for 2 hours at room temperature before centrifuging for 20 min at 2000 x g. Collected serum was snap-frozen in liquid nitrogen and kept in -80°C. ELISA kits (mouse CRP (MCRP00), mouse IL-6 (M6000B), mouse CCL2/MCP-1 (MJE00B)) were purchased from R&D Systems and performed following the manufacturer’s instruction.

#### Dual-energy X-ray absorptiometry (DEXA) scanning

Euthanized mice were scanned using iNSiGHT DEXA scanner (OsteoSys) at low energy (60 kV) and high energy (80 kV), 0.80 mA current, for 5 seconds of scanning at each energy level. An embedded software was used to compile the images and calculate the mass and area of fat.

#### Tissue collection and digestion

For synovial tissue collection and digestion, skin and surrounding muscle above the knee joint was excised, patellar ligament cut with scissors, and surrounding soft tissue micro-dissected away and placed into digestion media consisting of DMEM/F12 (Corning) with 10% Fetal Bovine Serum (FBS; Corning), 1% penicillin/streptomycin/amphotericin B solution (P/S/A; Corning), 1 mg/mL dispase (Gibco), 1 mg/mL type 2 collagenase (Worthington), 10 µg/mL gentamycin (Teknova) and 100 μg/mL primocin (Invivogen). The infrapatellar fat pad and meniscus was also included, and digestion was performed at 37°C for 2-4 hours. For adipose tissue collection and digestion, visceral, inguinal, and epididymal fat pads were collected from HFD mice, minced with scissors, and digested in digestion media for 2-4 hours. For mouse femoral chondrocyte and pig articular chondrocyte collection and digestion, we used methods previously described by our group (75). All cells were filtered and centrifuged for 300xg for 10 minutes and used for further analyses.

For bone marrow harvest, two methods were used based on needed quantities. For a smaller quantity of bone marrow, cells were flushed from the cut diaphysis of femurs and tibias adapted from previously described methods (76) using 1-2% FBS+PBS in a 27-gauge syringe and cells passed through a 70 μM cell filter. Bone marrow cells were centrifuged at 300xg for 10 minutes and red blood cells were lysed on ice with Ammonium Chloride solution (STEMCELL Tech #07850). For a larger quantity of bone marrow, long bones from mice were harvested and crushed with a mortar and pestle as previously described (77), filtered, and passed through a Ficoll-Paque density gradient (Sigma #GE17-1440-02) to remove bone fragments and isolate the polymorphonuclear cells. Red blood cells were lysed as described and bone marrow cells were used for further analyses.

#### Histology and immunocytochemistry

Mouse livers and adipose tissues were dissected and fixed in 4% PFA for 24 hour at 4 °C, where mouse legs were further decalcified with 14% EDTA (pH 7.4) for 14-21 days at 4 °C before paraffin embedding. Tissues were embedded in paraffin and cut at 5 µm thick. Antigen was retrieved by Tris HCL (pH 10; Sigma) at 95°C. Sections were blocked with 2.5% normal horse serum for 1 hour at room temperature and incubated with primary antibody (CD68, Invitrogen, PA5-178996) in 1% BSA at 4 °C overnight. Slides were washed and incubated at room temperature for 1 hour in secondary antibody-HRP (Vector Laboratories, MP-7401). Antibodies were then visualized by peroxidase substrate kit DAB (Vector Laboratories, SK-4100). For Picrosirius Red staining, slides were stained with Picro-Sirius Red Stain Kit (Abcam, ab15068) according to the manufacturer’s instructions.

For *in vivo* hind leg TRAP staining, deparaffinized and rehydrated sections were incubated in 50 mL TRAP staining medium (110 mM Sodium Acetate (Sigma, S-2889), 50 mM Tartaric Acid (Sigma, T-6521), 4.5 mM Naphthol AS-BI Phosphate (Sigma, N-2125) in distilled water for 1 hr at 37°C. After incubation, 1 mL of Pararosaniline Dye in Sodium Nitrite solution was added and incubated for 30 min at 37°C. Sections were counter-stained with 0.08% Fast Green for 90 sec. Toluidine Blue staining was performed in ordinance with general lab protocols and Research Society International (OARSI) scoring was performed as described previously (78). Slides were imaged using a Zeiss Axio Imager.A2 Microscope Axiocam 105 color camera with Zen 2 program at three random locations. Positive stain was quantified using ImageJ and an average of three images per sample was used for analysis.

For *in vitro* osteoclast TRAP staining, differentiated osteoclasts were fixed in 4% PFA at room temperature for 10-20 minutes and stained according to the Acid Phosphatase, Leukocyte (TRAP) Kit protocol (Sigma #387A). For immunocytochemistry, mouse chondrocytes were cultured on glass chamber slides (Falcon) and fixed with 4% PFA for 10-15 minutes. Fixed cells were washed with HBSS before cell permeabilization with 0.1% TritonX + HBSS, washed again, then blocked with 2% serum + HBSS for 1 hour. Cells were incubated with primary antibodies (gp130 - R&D cat#AF468; LAMP1 – Santa Cruz Biotech cat#sc-20011) diluted in 0.1% serum + HBSS overnight at 4 °C, washed after, then incubated with secondary antibodies for 1 hour at room temperature. Cells were counterstained with DAPI, mounted, and imaged with a Zeiss Axio Imager.A2 Microscope Axiocam 503 monochrome camera with the Zen.2 software.

#### Micro-computed tomography (μCT) and analysis

Fixed mouse legs were dissected and fixed in 4% PFA for 24 hours and scanned with XT H 225 ST (Nikon) in 20 µm resolution, at 80 kV energy, 120 μA current, and 9.6 W power at the USC Molecular Imaging Center. The trabecular bone under the growth plate near the proximal tibia was reconstructed and analyzed using Amira Software. 1 mm length of tibia bone mineral density was measured by bone volume over total volume (BV/TV).

#### RNA extraction, quantitative RT-PCR, RNA sequencing library preparation and analysis

RNA was extracted using the RNeasy Mini Kit (Qiagen) following the manufacturer’s protocol. cDNA was generated using the Maxima First Strand cDNA Synthesis Kit (Thermo Scientific) and Power SYBR Green (Applied Biosystems) was used for RT-PCR amplification. Detection was performed using the Step One Plus Real-Time PCR system (Applied Biosystems) and the comparative Ct method for relative quantification (1/2^(ΔCt)) was used to quantitate gene expression, where results were normalized to *Rpl7* (ribosomal protein L7) expression levels. In *in vitro* experiments with a control group, the dd-C_t_ (ΔΔC_t_) comparative method was used to normalize gene expression to the control group.

For bulk RNA sequencing, total RNA was quantified using a Qubit fluorometer and RNA quality was assessed using an Agilent Bioanalyzer 2100. Universal Plus mRNA-Seq Library with NuQuant (TECAN) was used to generate stranded RNA-seq libraries. Briefly, poly(A) RNA was selected followed by RNA fragmentation and double stranded cDNA was generated using a mixture of random and oilgo(dT) priming. Libraries were constructed by end repairing the cDNA to generate blunt ends, ligation of Unique Dual Index (UDI) adaptors, strand selection, and PCR amplification. Different adapters were used for multiplexing samples in one lane. Libraries were pooled and sequencing was performed at UCLA Technology Center for Genomics & Bioinformatics (TCGB) core on a NovaSeq SP/ X Plus with paired end 50 bp reads. Data quality check was done on Illumina SAV and demultiplexing was performed with Illumina CASAVA 1.8.2.

For bulk RNA sequencing analysis, the quality of raw fastq files were determined using FastQC (v0.11.9) (http://www.bioinformatics.babraham.ac.uk/projects/fastqc). Adapters were trimmed using Cutadapt (79); trimmed / filtered reads were aligned to mouse reference genome (mm39- GENCODE Release M30) using STAR aligner (version 2.7.8a) (80). Transcript levels were quantified to the reference using a modified expectation-maximization (EM) algorithm. Normalization was done using median ratio method and differential expression analysis was performed using DESeq2 (v3.5) (81), where genes were considered to be differentially expressed based on fold change > 1.5 and p-value < 0.05. Gene ontology enrichment analysis for differentially expressed genes was performed using Metascape (82), where genes were considered differentially expressed based on fold change > 0.05 and adjusted p-value < 0.01.

**Table 1.**
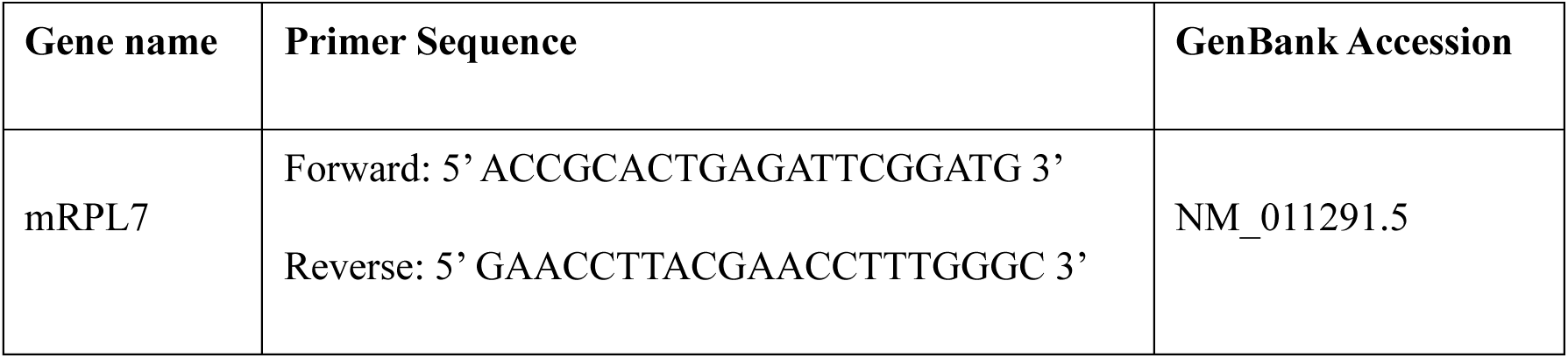
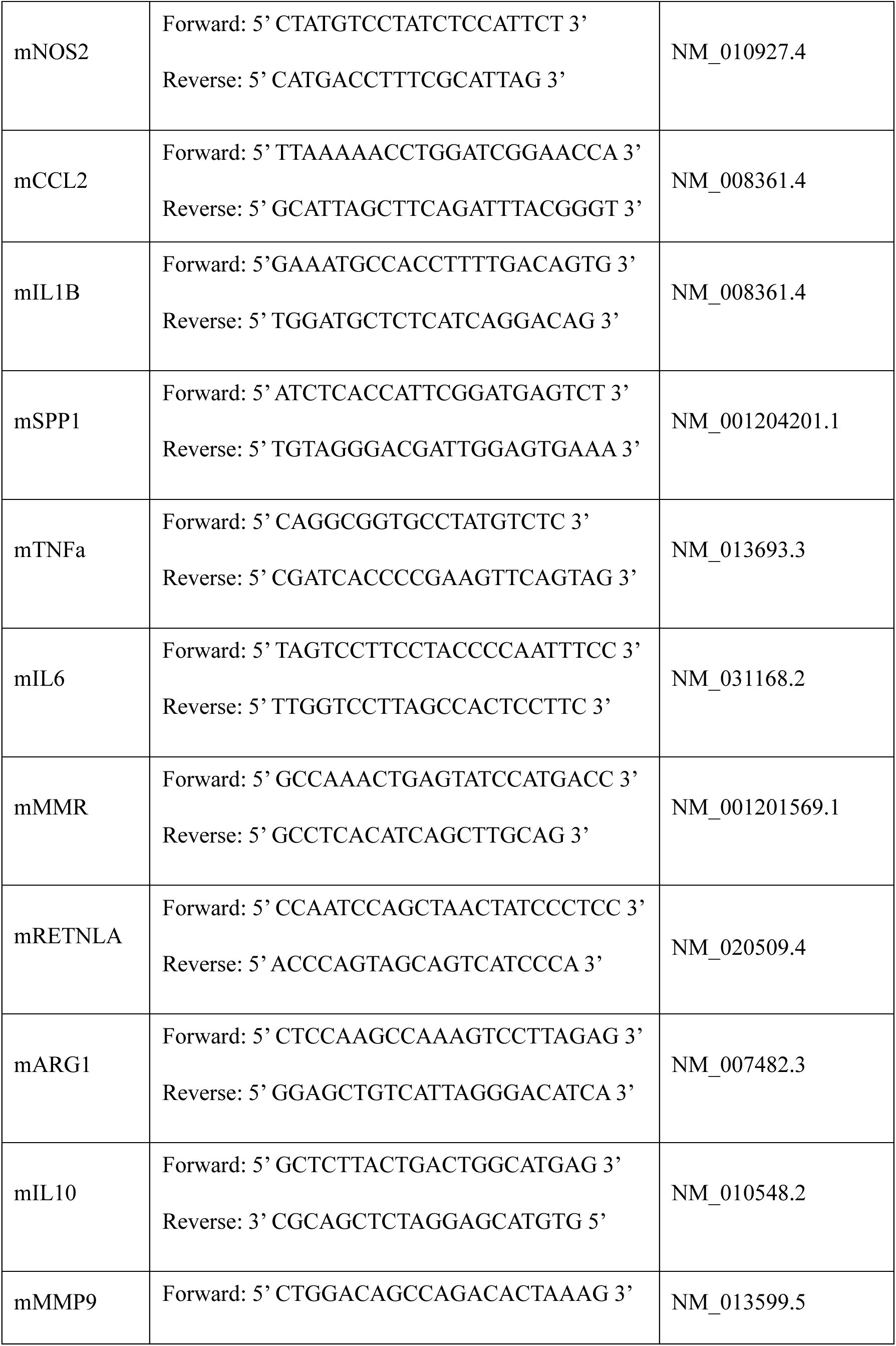
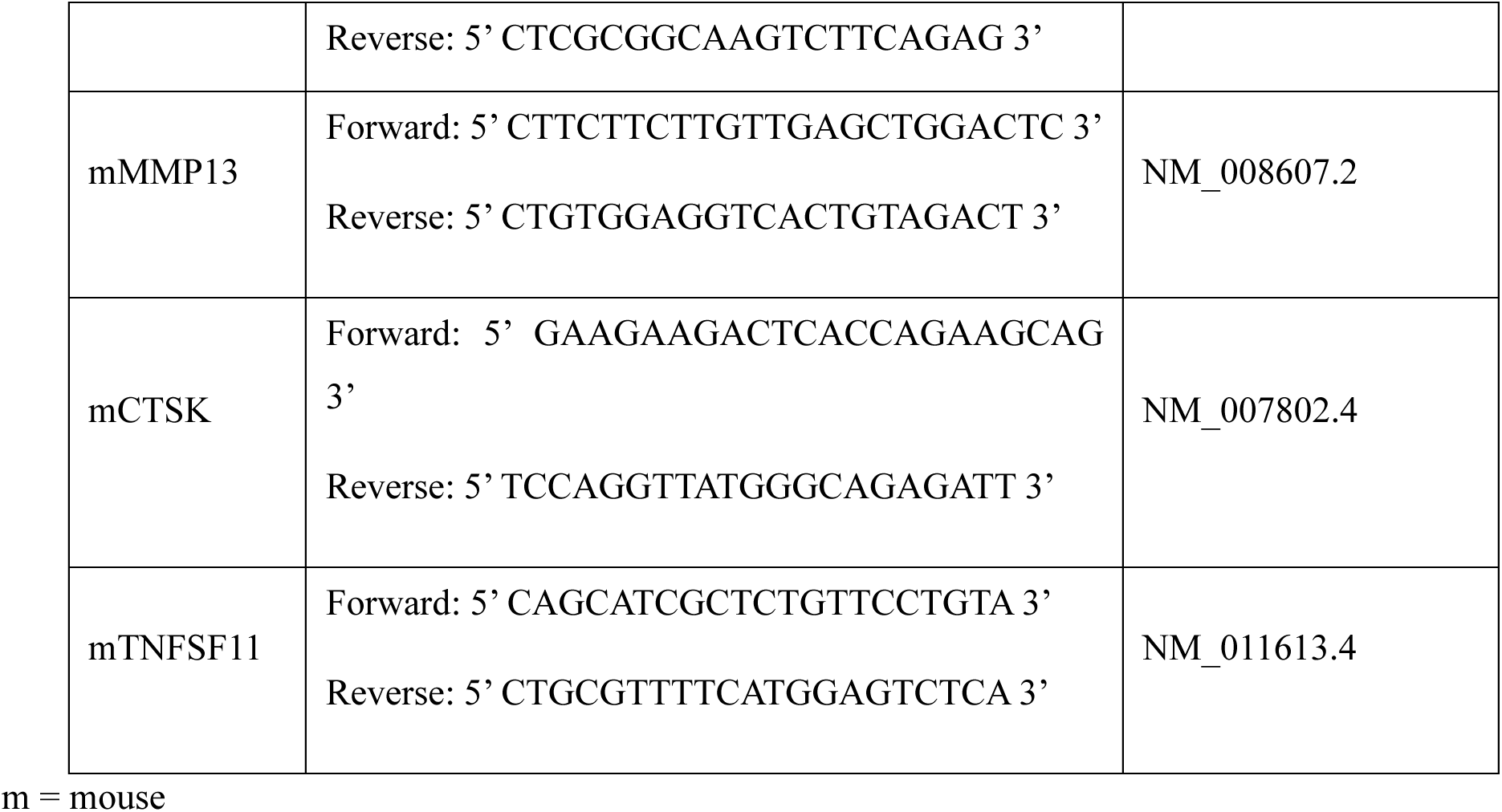
qPCR primer sequences.

#### Flow Cytometry and FACS

Flow cytometry and cell sorting were performed on a BD FACS Aria II and BD Symphony cell sorter. Cells were stained on ice for 20-30 minutes at a general concentration of 1 μL each antibody per 1x10^6^ cells in 1-2% FBS+PBS and viability was determined by exclusion of either DAPI (Fisher Scientific# 5748) or Live/Dead Aqua Viability Dye (Thermo #L34957). Flow cytometry data was analyzed using FlowJo software, and the anti-mouse antibodies used for this study were: Ly6G-BV605 (Biolegend #127639), Ly6G-PE (Biolegend #127607), CD11b-PE/Cy7 (Biolegend #101216), Ly6C-FITC (Biolegend #128006), CD45-APCFire750 (Biolegend #103154), F4/80-BV421 (Biolegend #123131), and gp130 (R&D #FAB4681A).

The Bromodeoxyuridine (BrdU) cell proliferation assay was performed according to the manufacturer’s protocols (Invitrogen #8812-6600 and BD #550891). Mice were injected intraperitoneally with 2 mg of BrdU (10 mg/ml stock) and peripheral blood and bone marrow was assessed 4 hours post injection. For the gp130 internalization assay measured by Flow cytometry, cells were stimulated with hyper IL6 for 30 or 60 minutes, stained, and fixed in 4% PFA for 10 minutes to retain cell surface-labeled gp130.

#### Cell culture

To derive bone marrow-derived macrophages *in vitro*, we followed previously a described macrophage differentiation protocol (83). Briefly, total bone marrow was cultured in 10% FBS + 1% PSA + DMEM/F12 supplemented with 50 ng/ml MCSF (Peprotech #315-02) for 5 days total, where fresh media was added to existing media on day 3. On day 5, formed macrophages were passaged into assay plates via trypsin and gentle scraping with a cell scraper. Bone marrow-derived macrophages were validated routinely via Flow Cytometry for mature macrophage-specific markers.

To generate osteoclasts *in vitro,* we followed a previously described protocol (84). Briefly, total bone marrow was plated in a cell culture flask with 10 ng/ml MCSF + 10% FBS + 1% PSA + αMEM for 24 hours. The non-adherent cells in the media were collected, centrifuged, and replated in media containing 30 ng/ml MCSF for 2 additional days. Then the cells were cultured in 30 ng/ml MCSF and 100 ng/ml RANKL (Thermo #315-11C) in media for 5 days total.

For mouse femoral and pig articular chondrocytes, passage 1-2 cells were maintained in 10% FBS + 1% PSA + DMEM/F12 and media was changed twice weekly. Generally, cells were serum starved either 2-4 hours (monocytes/ macrophages) or 16-24 hours (chondrocytes) before *in vitro* experiments to remove background signal from serum. Cytokines IL6 (Peprotech #200-06), hyper IL6 (Fisher #8954SR025), and OSM (Fisher #495-MO-025 or Thermo #300-10T-100) were used at final concentration of 20 ng/ml each.

#### Incucyte^TM^ live cell imaging

To image and measure gp130 endocytosis over time, the Incucyte^TM^ platform was used as previously described (85). BM-derived macrophages were plated in 96 well plates at 10,000 – 20,000 cells per well. An unconjugated anti-mouse gp130 antibody (Biolegend #149402) was labeled with a pH-sensitive dye following the manufacturer’s protocol (Sartorius #4737). Each condition had 3-4 technical replicate wells and images of each well were taken every 10 minutes for 8 hours on the Incucyte S3.

#### GTPase activation assay and Western blot

The Rac1/Cdc42 (Rho GTPases) activation assay was performed in ordinance with the manufacturer’s protocol (Cytoskeleton #BK034). Briefly, cell lysates were incubated with PAK-PBD affinity beads, which binds to the GTP-bound (active) pocket of Rac1 and Cdc42. Bound active Rac1/Cdc42 is eluted off the beads into SDS buffer and active Rac1/Cdc42 is analyzed via Western blot using anti-Rac1 and Cdc42 antibodies.

For Western blotting, cells were lysed in IP lysis buffer (Thermo #87788) supplemented with a phosphatase/protease inhibitor (Thermo #78441), clarified, and protein concentration was measured using the BCA assay (Thermo #23227). 1-10 μg total protein was mixed with Laemmli buffer (BioRad), boiled at 100°C for 5-6 minutes, and loaded into 4-15% SDS-PAGE TGX precast gels (BioRad). Gels were transferred via semi-dry Trans-Blot Turbo onto a 0.2 μM pore PVDF membrane (BioRad) and blocked with Everyblot Blocking Buffer (BioRad #12010020) for 10 minutes. Blocked membranes were incubated with primary antibodies (phospho-p38, total p38, and H3 at 1:1000 dilution; Cell Signaling Tech. #4511, #9212 and #9715 respectively) diluted in Everyblot buffer overnight at 4°C. After washing unbound primary antibodies off, membranes were incubated with anti-rabbit IgG-HRP secondary antibody (Thermo #31460) for 1 hour at room temperature. After washing, membranes were developed with Clarity Western ECL blotting substrate (BioRad #1705061) and imaged on the BioRad ChemiDoc Touch platform.

#### MMP activity assay

General MMP activity was measured following the manufacturer’s protocol (Anaspec #AS-71158). Briefly, passage 1-2 femoral chondrocytes were serum starved overnight and cytokines were added directly to the wells for 6 and 24 hours total. Cell culture media that contained secreted MMPs was collected at the given time points and analyzed using the SensoLyte fluorometric kit; cells were also lysed for qPCR analysis.

#### Small molecule inhibitor assay

Pig articular chondrocytes were used at passage 1-2, serum starved in 1% charcoal-stripped FBS + DMEM overnight, stimulated with OSM (20 ng/ml) and treated with either vehicle (DMSO), p38 inhibitor at 10 uM (SB203580-1, MedChemExpress), Erk 1/2 inhibitor at 12 nM (SCH772984-1, MedChemExpress), SFK inhibitor at 0.84 uM (SU6656-1, MedChemExpress), or Pi3K inhibitor at 3 uM (PI828-1, Tocris). All inhibitors and vehicle were added to 1% charcoal-stripped FBS + DMEM and cells were lysed for qPCR after 24 hours.

#### Plasmid isolation

Glycerol stocks for pCMV-VSV-G, (Plasmid #8454) pCMV-dR8.2 dvpr, (Plasmid #8455) pEGFP-N1-FLAG, (Plasmid #60360) and GFP-Dynamin 2 K44A, (Plasmid #22301) were obtained from Addgene. After streaking in Luria-Bertani agar plates, single colonies were picked and amplified for plasmid isolation. Plasmids were harvested following the QIAGEN Plasmid Maxi Kit protocol.

#### Viral Vector Generation and Transduction

pEGFP-N1-FLAG and GFP-Dynamin 2 K44A were cloned into a viral backbone with support from VectorBuilder. pCMV-VSV-G, pCMV-dR8.2 dvpr, along with either pEGFP-N1-FLAG or GFP-Dynamin 2 K44A (2:5:5) were transfected into HEK293T cells (ATCC) using antibiotic free DMEM/F12 (Corning) with 10% FBS (Corning) and LipoFexin following mannufacturer’s protocol. The media wsa changed to complete media after 12hrs to reduce cytotoxicity. Supernatant containing lentivirus was collected at 48hrs and 72hrs and filtered using 0.45um filter. Lentivirus containing supernatant was concentrated following the protocol of Lenti-X concentrator (Takara) and centrifuged. Concentrated lentivirus was resuspended in complete DMEM/F12 media and used to transduce pig chondrocytes alongwith polybrene (Sigma). All assays with the transduced cells were performed 48hrs after transduction.

## Data Availability

All data is deposited in GEO data base and is available under the accession numbers GSEXXXX.

## Acknowledgments

We would like to acknowledge the USC Stem Cell Flow Cytometry Core and Imaging Core for their continued support and assistance. Schematics were made using Biorender.com. Research in this study was supported by the Department of Defense grant W81XWH-13-1-0465 (DE), National Institute of Health grants R01AG058624, R61NS131307, R01AR071734, 1R01AR085642-01 (DE) and National Institute of Dental and Craniofacial Research grant T90DE021982 (JT).

## Author contributions

J.T. conceptualized the study, performed and analyzed experiments, wrote and edited the manuscript. Y.L. conceptualized the study, performed and analyzed experiments. A.S. performed bulk RNA sequencing data, wrote and edited the manuscript. J.Y, U.S., N.L., J.L., A.D., V. J. and M.V. processed tissues and performed experiments. A.M.M., L.G., and R.L. supervised experiments and edited the manuscript. D.E. conceptualized, supervised the study and edited the manuscript.

## References

1. Ferrero-Miliani L, Nielsen OH, Andersen PS, Girardin SE. Chronic inflammation: importance of NOD2 and NALP3 in interleukin-1beta generation. Clin Exp Immunol. 2007 Feb;147(2):227–35.

2. Furman D, Campisi J, Verdin E, Carrera-Bastos P, Targ S, Franceschi C, et al. Chronic inflammation in the etiology of disease across the life span. Nat Med. 2019 Dec;25(12):1822–32.

3. Chavda VP, Feehan J, Apostolopoulos V. Inflammation: The Cause of All Diseases. Cells. 2024 Nov 18;13(22):1906.

4. Menzel A, Samouda H, Dohet F, Loap S, Ellulu MS, Bohn T. Common and Novel Markers for Measuring Inflammation and Oxidative Stress Ex Vivo in Research and Clinical Practice-Which to Use Regarding Disease Outcomes? Antioxidants (Basel). 2021 Mar 9;10(3):414.

5. Bettcher BM, Neuhaus J, Wynn MJ, Elahi FM, Casaletto KB, Saloner R, et al. Increases in a Pro-inflammatory Chemokine, MCP-1, Are Related to Decreases in Memory Over Time. Front Aging Neurosci. 2019;11:25.

6. Nedunchezhiyan U, Varughese I, Sun AR, Wu X, Crawford R, Prasadam I. Obesity, Inflammation, and Immune System in Osteoarthritis. Front Immunol. 2022;13:907750.

7. Zheng L, Zhang Z, Sheng P, Mobasheri A. The role of metabolism in chondrocyte dysfunction and the progression of osteoarthritis. Ageing Res Rev. 2021 Mar;66:101249.

8. Alberro A, Iribarren-Lopez A, Sáenz-Cuesta M, Matheu A, Vergara I, Otaegui D. Inflammaging markers characteristic of advanced age show similar levels with frailty and dependency. Sci Rep. 2021 Feb 23;11(1):4358.

9. Khaodhiar L, Ling PR, Blackburn GL, Bistrian BR. Serum levels of interleukin-6 and C-reactive protein correlate with body mass index across the broad range of obesity. JPEN J Parenter Enteral Nutr. 2004;28(6):410–5.

10. Rose-John S. Interleukin-6 signalling in health and disease. F1000Res. 2020;9:F1000 Faculty Rev-1013.

11. Stahl N, Boulton TG, Farruggella T, Ip NY, Davis S, Witthuhn BA, et al. Association and activation of Jak-Tyk kinases by CNTF-LIF-OSM-IL-6 beta receptor components. Science. 1994 Jan 7;263(5143):92–5.

12. Fischer P, Hilfiker-Kleiner D. Role of gp130-mediated signalling pathways in the heart and its impact on potential therapeutic aspects. Br J Pharmacol. 2008 Mar;153 Suppl 1(Suppl 1):S414-427.

13. Taniguchi K, Wu LW, Grivennikov SI, de Jong PR, Lian I, Yu FX, et al. A gp130-Src-YAP module links inflammation to epithelial regeneration. Nature. 2015 Mar 5;519(7541):57–62.

14. Honke N, Ohl K, Wiener A, Bierwagen J, Peitz J, Di Fiore S, et al. The p38-mediated rapid down-regulation of cell surface gp130 expression impairs interleukin-6 signaling in the synovial fluid of juvenile idiopathic arthritis patients. Arthritis Rheumatol. 2014 Feb;66(2):470–8.

15. Wang W, Bian J, Sun Y, Li Z. The new fate of internalized membrane receptors: Internalized activation. Pharmacol Ther. 2022 May;233:108018.

16. Schmidt-Arras D, Rose-John S. Endosomes as Signaling Platforms for IL-6 Family Cytokine Receptors. Front Cell Dev Biol. 2021;9:688314.

17. German CL, Sauer BM, Howe CL. The STAT3 beacon: IL-6 recurrently activates STAT 3 from endosomal structures. Exp Cell Res. 2011 Aug 15;317(14):1955–69.

18. Honke N, Ohl K, Wiener A, Bierwagen J, Peitz J, Di Fiore S, et al. The p38-mediated rapid down-regulation of cell surface gp130 expression impairs interleukin-6 signaling in the synovial fluid of juvenile idiopathic arthritis patients. Arthritis Rheumatol. 2014 Feb;66(2):470–8.

19. Weller SG, Capitani M, Cao H, Micaroni M, Luini A, Sallese M, et al. Src kinase regulates the integrity and function of the Golgi apparatus via activation of dynamin 2. Proc Natl Acad Sci U S A. 2010 Mar 30;107(13):5863–8.

20. Hikita T, Kuwahara A, Watanabe R, Miyata M, Oneyama C. Src in endosomal membranes promotes exosome secretion and tumor progression. Sci Rep. 2019 Mar 1;9(1):3265.

21. Miao MZ, Su QP, Cui Y, Bahnson EM, Li G, Wang M, et al. Redox-active endosomes mediate α5β1 integrin signaling and promote chondrocyte matrix metalloproteinase production in osteoarthritis. Sci Signal. 2023 Oct 31;16(809):eadf8299.

22. Kim MJ, Byun JY, Yun CH, Park IC, Lee KH, Lee SJ. c-Src-p38 mitogen-activated protein kinase signaling is required for Akt activation in response to ionizing radiation. Mol Cancer Res. 2008 Dec;6(12):1872–80.

23. DerMardirossian C, Rocklin G, Seo JY, Bokoch GM. Phosphorylation of RhoGDI by Src regulates Rho GTPase binding and cytosol-membrane cycling. Mol Biol Cell. 2006 Nov;17(11):4760–8.

24. Shkhyan R, Flynn C, Lamoure E, Sarkar A, Van Handel B, Li J, et al. Inhibition of a signaling modality within the gp130 receptor enhances tissue regeneration and mitigates osteoarthritis. Sci Transl Med. 2023 Mar 22;15(688):eabq2395.

25. Bae HR, Shin SK, Yoo JH, Kim S, Young HA, Kwon EY. Chronic inflammation in high-fat diet-fed mice: Unveiling the early pathogenic connection between liver and adipose tissue. J Autoimmun. 2023 Sept;139:103091.

26. Kratofil RM, Kubes P, Deniset JF. Monocyte Conversion During Inflammation and Injury. Arterioscler Thromb Vasc Biol. 2017 Jan;37(1):35–42.

27. Hoenow S, Yan K, Noll J, Groneberg M, Casar C, Lory NC, et al. The Properties of Proinflammatory Ly6Chi Monocytes Are Differentially Shaped by Parasitic and Bacterial Liver Infections. Cells. 2022 Aug 16;11(16):2539.

28. Rose-John S. Interleukin-6 signalling in health and disease. F1000Res. 2020;9:F1000 Faculty Rev-1013.

29. Fischer M, Goldschmitt J, Peschel C, Brakenhoff JP, Kallen KJ, Wollmer A, et al. I. A bioactive designer cytokine for human hematopoietic progenitor cell expansion. Nat Biotechnol. 1997 Feb;15(2):142–5.

30. Torres-Guzman AM, Morado-Urbina CE, Alvarado-Vazquez PA, Acosta-Gonzalez RI, Chávez-Piña AE, Montiel-Ruiz RM, et al. Chronic oral or intraarticular administration of docosahexaenoic acid reduces nociception and knee edema and improves functional outcomes in a mouse model of Complete Freund’s Adjuvant-induced knee arthritis. Arthritis Res Ther. 2014 Mar 10;16(2):R64.

31. Kim JM, Lin C, Stavre Z, Greenblatt MB, Shim JH. Osteoblast-Osteoclast Communication and Bone Homeostasis. Cells. 2020 Sept 10;9(9):2073.

32. Heymann D, Rousselle AV. gp130 Cytokine family and bone cells. Cytokine. 2000 Oct;12(10):1455–68.

33. Sims NA. gp130 signaling in bone cell biology: multiple roles revealed by analysis of genetically altered mice. Mol Cell Endocrinol. 2009 Oct 30;310(1–2):30–9.

34. Sims NA, Jenkins BJ, Quinn JMW, Nakamura A, Glatt M, Gillespie MT, et al. Glycoprotein 130 regulates bone turnover and bone size by distinct downstream signaling pathways. J Clin Invest. 2004 Feb;113(3):379–89.

35. Shin HI, Divieti P, Sims NA, Kobayashi T, Miao D, Karaplis AC, et al. Gp130-mediated signaling is necessary for normal osteoblastic function in vivo and in vitro. Endocrinology. 2004 Mar;145(3):1376–85.

36. O’Brien CA, Gubrij I, Lin SC, Saylors RL, Manolagas SC. STAT3 activation in stromal/osteoblastic cells is required for induction of the receptor activator of NF-kappaB ligand and stimulation of osteoclastogenesis by gp130-utilizing cytokines or interleukin-1 but not 1,25-dihydroxyvitamin D3 or parathyroid hormone. J Biol Chem. 1999 July 2;274(27):19301–8.

37. Fujiwara Y, Piemontese M, Liu Y, Thostenson JD, Xiong J, O’Brien CA. RANKL (Receptor Activator of NFκB Ligand) Produced by Osteocytes Is Required for the Increase in B Cells and Bone Loss Caused by Estrogen Deficiency in Mice. J Biol Chem. 2016 Nov 25;291(48):24838–50.

38. Cao H, Garcia F, McNiven MA. Differential distribution of dynamin isoforms in mammalian cells. Mol Biol Cell. 1998 Sept;9(9):2595–609.

39. Ferguson SM, Brasnjo G, Hayashi M, Wölfel M, Collesi C, Giovedi S, et al. A selective activity-dependent requirement for dynamin 1 in synaptic vesicle endocytosis. Science. 2007 Apr 27;316(5824):570–4.

40. Cook TA, Urrutia R, McNiven MA. Identification of dynamin 2, an isoform ubiquitously expressed in rat tissues. Proc Natl Acad Sci U S A. 1994 Jan 18;91(2):644–8.

41. Bruzzaniti A, Neff L, Sanjay A, Horne WC, De Camilli P, Baron R. Dynamin forms a Src kinase-sensitive complex with Cbl and regulates podosomes and osteoclast activity. Mol Biol Cell. 2005 July;16(7):3301–13.

42. Weller SG, Capitani M, Cao H, Micaroni M, Luini A, Sallese M, et al. Src kinase regulates the integrity and function of the Golgi apparatus via activation of dynamin 2. Proc Natl Acad Sci U S A. 2010 Mar 30;107(13):5863–8.

43. Hikita T, Kuwahara A, Watanabe R, Miyata M, Oneyama C. Src in endosomal membranes promotes exosome secretion and tumor progression. Sci Rep. 2019 Mar 1;9(1):3265.

44. Ahn S, Kim J, Lucaveche CL, Reedy MC, Luttrell LM, Lefkowitz RJ, et al. Src-dependent tyrosine phosphorylation regulates dynamin self-assembly and ligand-induced endocytosis of the epidermal growth factor receptor. J Biol Chem. 2002 July 19;277(29):26642–51.

45. Lehtola T, Nummenmaa E, Tuure L, Hämäläinen M, Nieminen RM, Moilanen T, et al. Dexamethasone Attenuates the Expression of MMP-13 in Chondrocytes through MKP-1. Int J Mol Sci. 2022 Mar 31;23(7):3880.

46. Ravanti L, Heino J, López-Otín C, Kähäri VM. Induction of collagenase-3 (MMP-13) expression in human skin fibroblasts by three-dimensional collagen is mediated by p38 mitogen-activated protein kinase. J Biol Chem. 1999 Jan 22;274(4):2446–55.

47. Honke N, Ohl K, Wiener A, Bierwagen J, Peitz J, Di Fiore S, et al. The p38-mediated rapid down-regulation of cell surface gp130 expression impairs interleukin-6 signaling in the synovial fluid of juvenile idiopathic arthritis patients. Arthritis Rheumatol. 2014 Feb;66(2):470–8.

48. Li Z, Dai A, Yang M, Chen S, Deng Z, Li L. p38MAPK Signaling Pathway in Osteoarthritis: Pathological and Therapeutic Aspects. J Inflamm Res. 2022;15:723–34.

49. Weller SG, Capitani M, Cao H, Micaroni M, Luini A, Sallese M, et al. Src kinase regulates the integrity and function of the Golgi apparatus via activation of dynamin 2. Proc Natl Acad Sci U S A. 2010 Mar 30;107(13):5863–8.

50. Hikita T, Kuwahara A, Watanabe R, Miyata M, Oneyama C. Src in endosomal membranes promotes exosome secretion and tumor progression. Sci Rep. 2019 Mar 1;9(1):3265.

51. Miao MZ, Su QP, Cui Y, Bahnson EM, Li G, Wang M, et al. Redox-active endosomes mediate α5β1 integrin signaling and promote chondrocyte matrix metalloproteinase production in osteoarthritis. Sci Signal. 2023 Oct 31;16(809):eadf8299.

52. Murakami M, Kamimura D, Hirano T. Pleiotropy and Specificity: Insights from the Interleukin 6 Family of Cytokines. Immunity. 2019 Apr 16;50(4):812–31.

53. Liu NQ, Lin Y, Li L, Lu J, Geng D, Zhang J, et al. gp130/STAT3 signaling is required for homeostatic proliferation and anabolism in postnatal growth plate and articular chondrocytes. Commun Biol. 2022 Jan 17;5(1):64.

54. Febbraio MA. gp130 receptor ligands as potential therapeutic targets for obesity. J Clin Invest. 2007 Apr;117(4):841–9.

55. Wallenius V, Wallenius K, Ahrén B, Rudling M, Carlsten H, Dickson SL, et al. Interleukin-6-deficient mice develop mature-onset obesity. Nat Med. 2002 Jan;8(1):75–9.

56. Ridker PM. From C-Reactive Protein to Interleukin-6 to Interleukin-1: Moving Upstream To Identify Novel Targets for Atheroprotection. Circ Res. 2016 Jan 8;118(1):145–56.

57. Hosaka K, Rojas K, Fazal HZ, Schneider MB, Shores J, Federico V, et al. Monocyte Chemotactic Protein-1-Interleukin-6-Osteopontin Pathway of Intra-Aneurysmal Tissue Healing. Stroke. 2017 Apr;48(4):1052–60.

58. Bae HR, Shin SK, Yoo JH, Kim S, Young HA, Kwon EY. Chronic inflammation in high-fat diet-fed mice: Unveiling the early pathogenic connection between liver and adipose tissue. J Autoimmun. 2023 Sept;139:103091.

59. MacParland SA, Liu JC, Ma XZ, Innes BT, Bartczak AM, Gage BK, et al. Single cell RNA sequencing of human liver reveals distinct intrahepatic macrophage populations. Nat Commun. 2018 Oct 22;9(1):4383.

60. Kanda H, Tateya S, Tamori Y, Kotani K, Hiasa K ichi, Kitazawa R, et al. MCP-1 contributes to macrophage infiltration into adipose tissue, insulin resistance, and hepatic steatosis in obesity. J Clin Invest. 2006 June;116(6):1494–505.

61. Wilkinson AN, Gartlan KH, Kelly G, Samson LD, Olver SD, Avery J, et al. Granulocytes Are Unresponsive to IL-6 Due to an Absence of gp130. J Immunol. 2018 May 15;200(10):3547–55.

62. Chen YH, van Zon S, Adams A, Schmidt-Arras D, Laurence ADJ, Uhlig HH. The Human GP130 Cytokine Receptor and Its Expression-an Atlas and Functional Taxonomy of Genetic Variants. J Clin Immunol. 2023 Dec 22;44(1):30.

63. Barksby HE, Hui W, Wappler I, Peters HH, Milner JM, Richards CD, et al. Interleukin-1 in combination with oncostatin M up-regulates multiple genes in chondrocytes: implications for cartilage destruction and repair. Arthritis Rheum. 2006 Feb;54(2):540–50.

64. Li Z, Dai A, Yang M, Chen S, Deng Z, Li L. p38MAPK Signaling Pathway in Osteoarthritis: Pathological and Therapeutic Aspects. J Inflamm Res. 2022;15:723–34.

65. Zhu EY, Riordan JD, Vanneste M, Henry MD, Stipp CS, Dupuy AJ. SRC-RAC1 signaling drives drug resistance to BRAF inhibition in de-differentiated cutaneous melanomas. NPJ Precis Oncol. 2022 Oct 21;6(1):74.

66. Cao H, Yang P, Liu J, Shao Y, Li H, Lai P, et al. MYL3 protects chondrocytes from senescence by inhibiting clathrin-mediated endocytosis and activating of Notch signaling. Nat Commun. 2023 Oct 4;14(1):6190.

67. Clemente LP, Rabenau M, Tang S, Stanka J, Cors E, Stroh J, et al. Dynasore Blocks Ferroptosis through Combined Modulation of Iron Uptake and Inhibition of Mitochondrial Respiration. Cells. 2020 Oct 9;9(10):2259.

68. Preta G, Cronin JG, Sheldon IM. Dynasore - not just a dynamin inhibitor. Cell Commun Signal. 2015 Apr 10;13:24.

69. Weller SG, Capitani M, Cao H, Micaroni M, Luini A, Sallese M, et al. Src kinase regulates the integrity and function of the Golgi apparatus via activation of dynamin 2. Proc Natl Acad Sci U S A. 2010 Mar 30;107(13):5863–8.

70. Fessart D, Simaan M, Laporte SA. c-Src regulates clathrin adapter protein 2 interaction with beta-arrestin and the angiotensin II type 1 receptor during clathrin- mediated internalization. Mol Endocrinol. 2005 Feb;19(2):491–503.

71. Fessart D, Simaan M, Laporte SA. c-Src regulates clathrin adapter protein 2 interaction with beta-arrestin and the angiotensin II type 1 receptor during clathrin- mediated internalization. Mol Endocrinol. 2005 Feb;19(2):491–503.

72. Van Dyck PK, Piszkin L, Gorski EA, Nascimento ET, Abebe JA, Hoffmann LM, et al. Ionizable networks mediate pH-dependent allostery in the SH2 domain-containing signaling proteins SHP2 and SRC. Sci Signal. 2025 Nov 11;18(912):eadt3018.

73. Jiang D, Chowdhury AY, Nogalska A, Contreras J, Lee Y, Vergel-Rodriguez M, et al. Quantitative association between gene expression and blood cell production of individual hematopoietic stem cells in mice. Sci Adv. 2024 Jan 26;10(4):eadk2132.

74. Nees TA, Wang N, Adamek P, Zeitzschel N, Verkest C, La Porta C, et al. Role of TMEM100 in mechanically insensitive nociceptor un-silencing. Nat Commun. 2023 Apr 5;14(1):1899.

75. Tassey J, Sarkar A, Van Handel B, Lu J, Lee S, Evseenko D. A Single-Cell Culture System for Dissecting Microenvironmental Signaling in Development and Disease of Cartilage Tissue. Front Cell Dev Biol. 2021;9:725854.

76. Liu X, Quan N. Immune Cell Isolation from Mouse Femur Bone Marrow. Bio Protoc. 2015 Oct 20;5(20):e1631.

77. Jiang D, Chowdhury AY, Nogalska A, Contreras J, Lee Y, Vergel-Rodriguez M, et al. Quantitative association between gene expression and blood cell production of individual hematopoietic stem cells in mice. Sci Adv. 2024 Jan 26;10(4):eadk2132.

78. Lee Y, Tassey J, Sarkar A, Levi JN, Lee S, Liu NQ, et al. Pharmacological inactivation of a non-canonical gp130 signaling arm attenuates chronic systemic inflammation and multimorbidity induced by a high-fat diet. Sci Rep. 2024 Dec 28;14(1):31151.

79. Martin M. Cutadapt removes adapter sequences from high-throughput sequencing reads. EMBnet.journal. 2011;17(1):10–2.

80. Dobin A, Davis CA, Schlesinger F, Drenkow J, Zaleski C, Jha S, et al. STAR: ultrafast universal RNA-seq aligner. Bioinformatics. 2013 Jan 1;29(1):15–21.

81. Love MI, Huber W, Anders S. Moderated estimation of fold change and dispersion for RNA-seq data with DESeq2. Genome Biol. 2014;15(12):550.

82. Zhou Y, Zhou B, Pache L, Chang M, Khodabakhshi AH, Tanaseichuk O, et al. Metascape provides a biologist-oriented resource for the analysis of systems-level datasets. Nat Commun. 2019 Apr 3;10(1):1523.

83. Haag SM, Murthy A. Murine Monocyte and Macrophage Culture. Bio Protoc. 2021 Mar 20;11(6):e3928.

84. Chevalier C, Çolakoğlu M, Brun J, Thouverey C, Bonnet N, Ferrari S, et al. Primary mouse osteoblast and osteoclast culturing and analysis. STAR Protoc. 2021 June 18;2(2):100452.

85. Maretti-Mira AC, Golden-Mason L, Salomon MP, Kaplan MJ, Rosen HR. Cholesterol-Induced M4-Like Macrophages Recruit Neutrophils and Induce NETosis. Front Immunol. 2021;12:671073.

